# 4D spatial transcriptomics reveals nodule identity emerges through stacked parallel developmental programs

**DOI:** 10.64898/2026.06.23.733685

**Authors:** Min-Yao Jhu, Jo Heffer, Chongjing Xia, Thiago Alexandre Moraes, Claire M. Mulvey, Dario Bressan, Chen Liu, Jin-Peng Gao, Philip Poole, Katharina Schiessl, Giles E. D. Oldroyd

**Author notes:** Senior author (Dario Bressan, Philip Poole, Katharina Schiessl, Giles E. D. Oldroyd). Lead contacts (Min-Yao Jhu).

## Abstract

Legume root nodules enable symbiotic nitrogen fixation through the development of specialized cells that accommodate nitrogen-fixing bacteria intracellularly and support bacterial nitrogenase activity. Here, we present a 4D (3D space and time) spatial transcriptomic atlas of *Medicago truncatula* nodules and lateral roots, revealing specialized symbiotic cell types that develop alongside a conserved lateral-root-derived program that underpins vascularization. Spatial profiling of both plant and bacterial transcripts resolves distinct transcriptional states and previously unrecognized cell states. Spatial analysis of developmental regulator mutants uncovers a cascading series of cell-type-specific programs during nodule maturation. *LSH1/LSH2* are central regulators of these programs, and *lsh1/lsh2* mutants exhibit collapse of hormonal gradients and nodule identity. Strikingly, loss of nodule identity collapses to a primary-root identity rather than lateral-root fate. This work reveals how tissue complexity emerges through stacked developmental programs sustained in distinct cellular compartments, allowing the emergence of cell types specialized for harboring nitrogen-fixing bacteria.

**HIGHLIGHTS:**

- >136,000 cells define a 4D spatial atlas of nodules and lateral roots
- 3D spatial and dual-species analysis resolves dynamic host and rhizobial cell states
- A shared meristem generates 3 spatially coordinated symbiotic, non-symbiotic, and vascular cell programs
- *LSH1/LSH2* are critical for regulatory programs underlying nodule development and identity
- Nodule identity loss collapses toward a primary root-associated state, not a lateral root fate

## INTRODUCTION

Symbiotic nitrogen fixation in legumes occurs in nodules: specialized organs, principally located on roots, that facilitate microbial colonization and nitrogen fixation in a controlled cellular environment^1,2^. In *Medicago truncatula*, indeterminate nodules maintain a persistent meristem and longitudinal zonation, including infection zone, interzone, nitrogen-fixation and senescence zones^3–6^. Genetic studies have defined many of the core signaling and transcriptional nodes that are required for nodulation^2,3,7–9^. Nodule organogenesis is widely viewed as an evolutionary co-option of the lateral root developmental program^1,10–12^. However, mature nodules acquire highly specialized tissue organization and symbiotic cell identities that are absent from lateral roots, raising a central unresolved question: how are conserved root developmental programs rewired to establish and maintain nodule identity during organogenesis? More specifically, when and where do distinct nodule cell identities emerge during primordium development, how are they coordinated with symbiotic bacterial states, and what developmental states arise when nodule identity regulators fail?

Nodule organogenesis is controlled by a regulatory network that integrates symbiosis signaling with conserved root developmental programs. Central to nodulation is the transcription factor *NODULE INCEPTION* (*NIN*)^13–16^, which operates downstream of symbiosis and cytokinin signalling^13^ mediated by the cytokinin receptor *CRE1*^17,18^. *NIN* activates both nodulation-specific and lateral-root-associated developmental regulators, including the canonical lateral-root regulator *Lateral Organ Boundaries Domain 16* (*LBD16*) ^10,19^, and core root meristem regulators, like *PLETHORAs* (*PLTs*)^20^, *WUSCHEL-related homeobox 5* (*WOX5*), and the *SHORTROOT–SCARECROW* (*SHR–SCR*)^21^ module, reflecting shared developmental programs between nodules and lateral roots. However, nodule development also requires organ-identity regulators, including *NF-YA1*^10,14^ and the *LSH1/LSH2–NOOT1/NOOT2* module^2,22,23^, together with coordinated auxin, cytokinin, and gibberellin signalling^7,24,25^. These studies suggest that nodules arise through rewiring of root developmental programs, but how these regulatory programs are spatiotemporally coordinated and established at cellular resolution remains poorly understood.

Recent single-cell and single-nucleus transcriptomic atlases have resolved transcriptional heterogeneity masked in prior bulk RNA-seq approaches^26–29^. Early-timepoint datasets revealed coordinated epidermal and internal tissue responses to rhizobia and cell-type specificity of nodulation-associated signalling^26,28^. Mature-nodule single-cell atlases clarified the functional division of labor within the nitrogen-fixation zone and provided inferred trajectories linking meristematic states to infected and uninfected fates^27^. However, protoplast or nuclear dissociation can introduce cellular and transcriptional biases^30,31^, underrepresent rare or fragile or structurally embedded populations^31^, and remove the spatial context^32,33^. Therefore, spatial relationships and lineage progression must be inferred indirectly from marker genes and transcriptional similarity. This limitation is particularly important in plants, where gene duplication and functional divergence can alter spatial expression domains even among conserved orthologs^34,35^, complicating marker transfer across species and developmental contexts. Consequently, mixed or unresolved clusters remain a recurring feature of current single-cell atlases. A spatially resolved developmental framework is therefore needed to determine when nodule cell identities emerge and how they are spatially organized within their native tissue context.

Here, we use Xenium *in situ*^36^ to construct a 4-dimensional (3-dimensional space + time) spatial transcriptomic atlas of nodules integrating subcellular-resolution gene expression with a developmental time-course analysis. By profiling 405 Medicago nodulation-associated genes across nodule development and 50 rhizobial genes in mature nodules, we generate a spatial atlas of nodule formation, maturation, and host–microbe transcriptional states. This framework resolves when and where transcriptionally similar precursor cells diverge toward distinct nodule- and root-associated fates, and how multiple developmental programs are coordinated within a shared primordium. Targeted genetic perturbations (*cre1, nf-ya1, lsh1/lsh2, noot1/noot2,* and *lbd16*) further reveal that disruption of key nodule identity regulators does not result in lateral-root conversion. Instead, identity-deficient nodules collapse toward a primary-root-associated transcriptional state that remains distinct from the canonical lateral-root developmental program. Together, this 4D spatial transcriptomic resource provides a framework for understanding how conserved root developmental modules are partitioned during indeterminate nodule and lateral-root organogenesis and generate the cellular complexity required for symbiotic nitrogen fixation.

## RESULTS

### Spatial anchoring resolves nodule transcriptomic cell identity

To establish a spatially grounded reference for mature nodule cell identity, we profiled wild-type *M. truncatula* nodules at 14 days post-inoculation (DPI) using the Xenium *in situ* spatial transcriptomics workflow^36^ with two panel designs: a 50-gene pilot and an expanded 405-gene panel (350-gene core plus 55-gene extension) (Figure 1). The panel captured hormone signaling components, cell-type markers, and representative lateral root and nodule developmental genes informed by published single-cell and single-nucleus^26,27^ and time-course bulk RNA-seq datasets from *M. truncatula*^10,23^.

**Figure 1.**
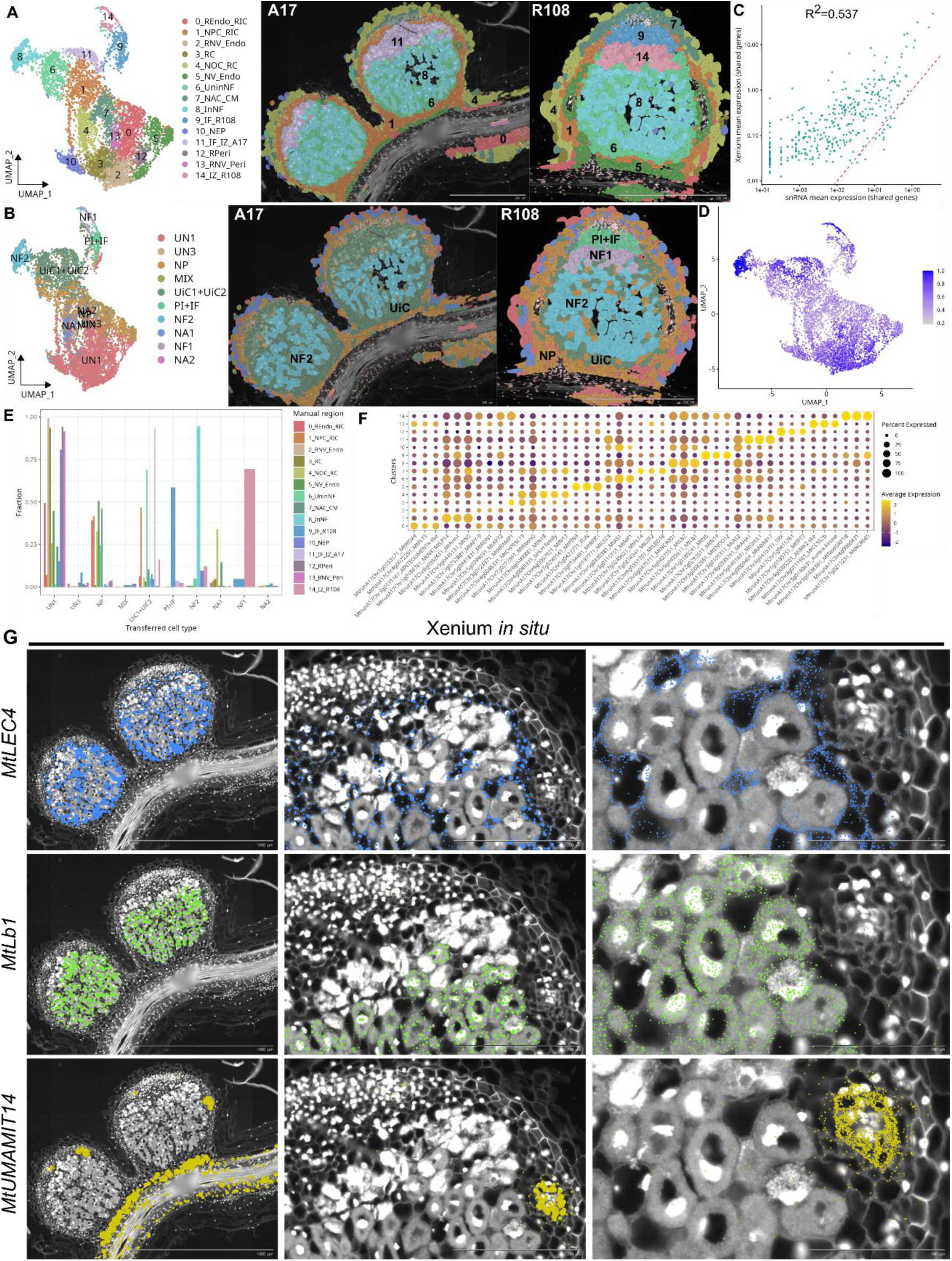
Spatial transcriptomics refines mature *Medicago* nodule cell-type annotatio. (A) Left, UMAP of the cell-wall–segmented spatial transcriptomic atlas of wild-type *Medicago truncatula* A17 and R108 nodules at 14 days post-inoculation (DPI), showing clustering based on spatial gene expression and histological context. Fifteen major cell types were identified: root endodermis and inner cortex (REndo_RIC, 0), nodule parenchyma cell and root inner cortex (NPC_RIC, 1), root and nodule vasculature (endodermis; RNV_Endo, 2), root cortex (RC, 3), nodule outer cortex and root cortex (NOC_RC, 4), nodule vasculature (endodermis; NV_Endo, 5), uninfected cells in the nitrogen fixation zone (UninNF, 6), nodule apex cortex and central meristem (NAC_CM, 7), infected cells in the nitrogen fixation zone (InNF, 8), infection zone specific to R108 (IF_R108, 9), nodule epidermis and parenchyma (NEP, 10), infection zone plus interzone in A17 (IF_IZ_A17, 11), root pericycle (RPeri, 12), root and nodule vasculature (pericycle; RNV_Peri, 13), and interzone specific to R108 (IZ_R108, 14). See Table SS3 for full cluster annotations. Right, spatial maps showing localisation of annotated cell types in representative A17 and R108 nodule sections. (B) UMAP and spatial maps of label transfer from a published 14-DPI single-cell RNA-seq atlas onto Xenium cells and tissue space. Abbreviations: NA1/NA2 (nodule apex), PI (pre-infection), IF (infection), NF1/NF2 (nitrogen fixation), NP (nodule parenchyma), VA (vascular), UiC1/UiC2 (uninfected cells), UN1–3 (unknown), MIX (mixed identity). (C) Normalized expression correlation between shared single-cell RNA-seq and Xenium genes. Each point represents one gene; the diagonal indicates equal expression. (D) UMAP colored by label-transfer confidence scores. (E) Comparison between spatially defined cell types and transferred single-cell labels. (F) Dot plot showing expression of marker genes defining spatial cell types identified in (A). (G) Xenium visualization of representative marker transcripts (*MtLEC4*, *Lb1*, *MtUMAMIT14*). See also Figure S1-2 and Table S1.

Cell wall-based segmentation enabled delineation of individual plant cells for quantitative spatial transcriptome analysis (Figure 1). Unsupervised clustering of spatial gene expression resolved 15 cell clusters corresponding to major anatomical domains, including infection zone, nitrogen-fixation zone, cortex, and vasculature, and these were largely conserved between wild-type *Medicago* A17 and R108 ecotypes (Figure 1A). Infection-zone and interzone populations were resolved separately in R108 (cluster 9 and 14), but remained merged in A17 (cluster 11), suggesting ecotype-specific zonation differences (Figure 1A). Spatial localization enabled anatomical annotation of transcriptionally similar cortical and vascular populations that are difficult to resolve in dissociation-based single-cell datasets (Figure 1A).

To benchmark these annotations, we performed label transfer from a published A17 14-DPI single-cell RNA-seq atlas^27^ to our A17 and R108 Xenium 14-DPI dataset (Figure 1B). Transferred cell labels were visualized both within the same UMAP embedding and mapped back onto the Xenium tissue sections (Figure 1B). While infection- and nitrogen-fixation-zone identities were largely recovered, cortical, vascular, and interzone populations showed substantial mixing and low-confidence assignments (Figure 1B, D). Notably, a large cluster previously annotated as “unknown 1^27^” in the single-cell atlas mapped onto multiple anatomically distinct spatial domains, including nodule cortex, root cortex, and vascular tissues, indicating that it represents a composite rather than a discrete cell identity (Figure 1B, E). Together, these analyses demonstrate that spatial context substantially improves resolution of transcriptionally similar nodule cell states and suggest that some single-cell-defined “cell types” represent mixtures of spatially distinct regulatory states, potentially reflecting both loss of spatial information and transcriptional changes associated with tissue dissociation.

Although Xenium profiles a targeted gene set rather than the whole transcriptome, it showed higher relative sensitivity for most shared genes after normalization, with moderate overall correspondence to single-cell RNA-seq (R² ≈ 0.537; Figure 1C). Label-transfer confidence was highest in transcriptionally distinct nitrogen-fixation-zone populations and lower in peripheral and vascular tissues (Figure 1D, E, S1), reinforcing the value of the spatial context for resolving anatomically adjacent cell states.

Using the spatially defined clusters, we identified the top 3 marker genes for all 15 cell clusters (Figure 1F), providing a spatially anchored marker resource for future cell-type identification and annotation. In plant biology, cell-type annotation often relies on marker genes identified through promoter-reporter lines, *in situ* hybridization, or orthologous markers transferred from model species such as *Arabidopsis*. However, promoter activity does not necessarily reflect endogenous transcript localization, and marker genes are not always conserved in their spatial expression domains across species. Although *in situ* hybridization can provide reliable spatial validation, it is labor-intensive and difficult to scale across large numbers of genes and developmental stages. By directly measuring transcript localization in intact tissues, spatial transcriptomics provides a higher resolution and systematic framework for establishing spatially informed marker gene resources. For instance, three genes previously used as single-cell markers^27^—*Lectin* (*MtLEC4*), *Leghemoglobin 1* (*MtLb1*), and *Usually Multiple Acids Move In and Out Transporters* (*MtUMAMIT14*) (Figure 1G), previously published using promoter GUS-based reporters^27^, in spatial transcriptomics revealed with much greater clarity of cellular resolution: *MtLEC4* associated with uninfected nitrogen-fixation cells, *MtLb1* to infected fixation-zone cells, and *MtUMAMIT14*, previously associated with an “unknown 1^27^” cluster, in spatial transcriptomics shown to be localized to vascular endodermal tissues, resolving previously ambiguous or unresolved annotations (Figure 1G).

Cell-wall and nucleus-anchored segmentation produced broadly concordant clusters and marker profiles, although each method showed different regional sensitivities (Figure S2, S3). Cell-wall segmentation better resolved enlarged cortical and fixation-zone cells, whereas parallel nucleus-anchored segmentation improved detection in densely packed meristematic and vascular tissues (Figure 1, S1). Thus, the segmentation strategy can affect regional sensitivity and can be optimized according to biological focus while preserving core cell-state structures.

### 3-dimensional spatial profiling captures late-stage mature nodule architecture and cell diversification

To fully capture late-stage structures that are difficult to resolve in single two-dimensional sections, including the peripheral vasculature (Figure S4) and sparsely distributed cell states, we constructed a 3D spatial transcriptomic atlas of 49-DPI nodules using serial Xenium profiling of paraffin sections (Figure 2), followed by histology-guided alignment (Figure S5A-B). In parallel, Serial Two-Photon Tomography (STPT) provided 3D visualization of whole nodule architecture and intrinsic autofluorescence (Figure S5C-D, Video S1). STPT resolved the 3D organization of the nodule peripheral vascular and autofluorescence from metabolically active rhizobia within the nitrogen-fixation zone (Figure S5C-D), which diminished toward the basal region, consistent with the presence of a senescence-associated zone characterized by decreased metabolic activity (Figure S5C-D, Video S1). This enabled reconstruction of late-stage nodule 3D phenotyping (Figure 2, Figure S5).

**Figure 2.**
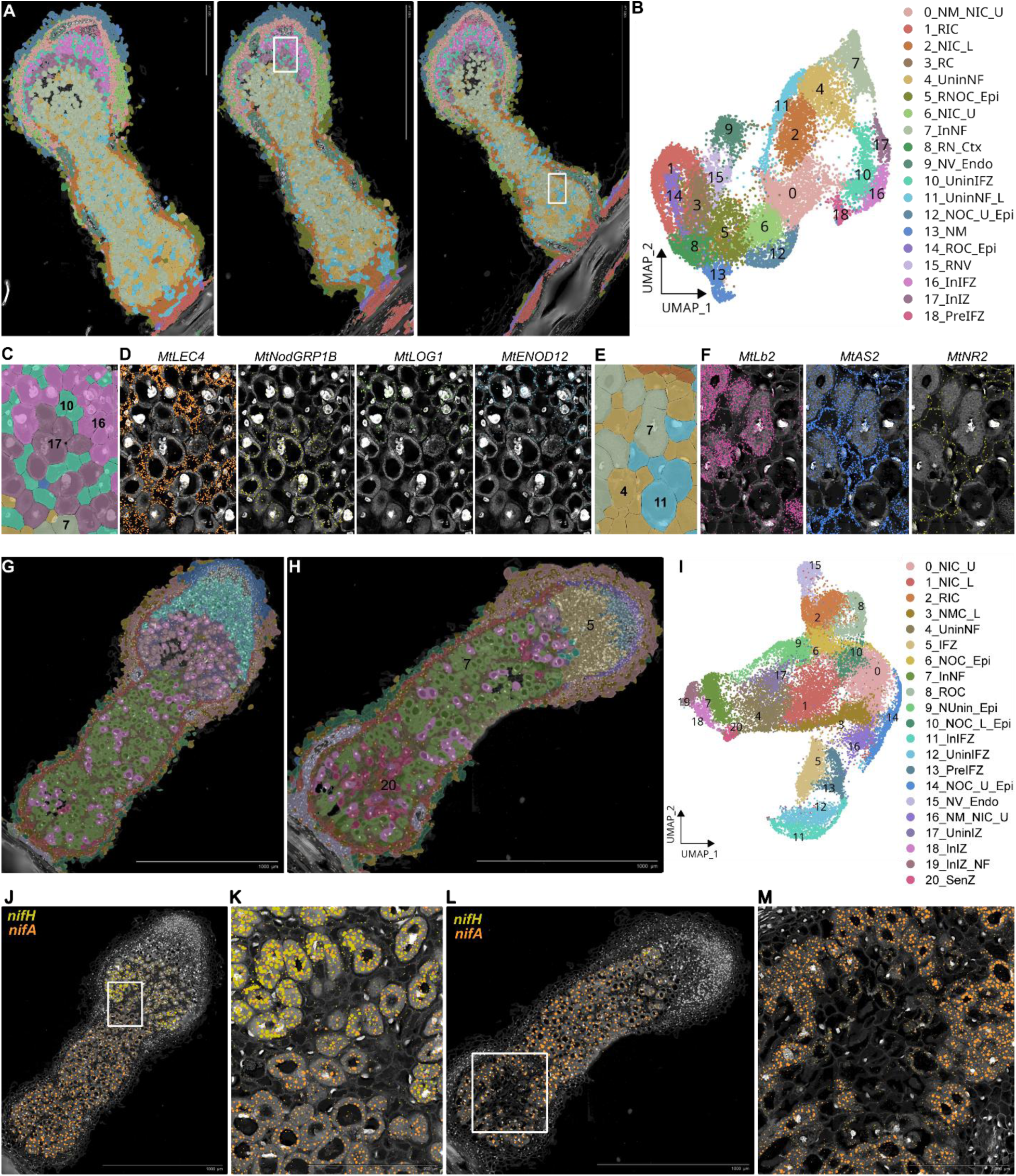
Dual-species 3D spatial transcriptomics resolves late-stage nodule architecture and rhizobial transcriptional states. (A–B) Representative Xenium sections and UMAP of the 49-DPI 3D spatial transcriptomic atlas. Nineteen clusters were resolved and annotated as follows: nodule meristem and upper nodule inner cortex (NM_NIC_U, cluster 0), root inner cortex (RIC, cluster 1), lower nodule inner cortex (NIC_L, cluster 2), root cortex (RC, cluster 3), uninfected cells of the nitrogen-fixation zone (UninNF, cluster 4), root and nodule outer cortex and epidermis (RNOC_Epi, cluster 5), upper nodule inner cortex (NIC_U, cluster 6), infected cells of the nitrogen-fixation zone (InNF, cluster 7), root and nodule cortex (RN_Ctx, cluster 8), nodule vasculature endodermis (NV_Endo, cluster 9), uninfected cells of the infection zone (UninIFZ, cluster 10), uninfected cells of the lower nitrogen-fixation zone (UninNF_L, cluster 11), upper nodule outer cortex and epidermis (NOC_U_Epi, cluster 12), nodule meristem (NM, cluster 13), root outer cortex and epidermis (ROC_Epi, cluster 14), root and nodule vasculature (RNV, cluster 15), infected cells of the infection zone (InIFZ, cluster 16), infected cells of the interzone (InIZ, cluster 17), and pre-infection zone cells (PreIFZ, cluster 18). (C–D) Spatial cell segregation and marker-gene expression of infected and uninfected populations in the infection zone and interzone. Infected cells (InIFZ, 16; InIZ, 17) are separated from uninfected infection-zone cells (UninIFZ, 10). Infection-associated populations express *MtNodGRP1B*, *MtLOG1*, and *MtENOD12*, whereas uninfected cells express *MtLEC4*. (E–F) Spatial cell segregation and marker-gene expression of nitrogen-fixation–zone populations. In addition to infected cells (InNF, 7), two uninfected populations are resolved: UninNF (4) and UninNF_L (11). Infected cells express *MtLb2*, whereas uninfected populations differentially express *MtAS2* and *MtNR2*. (G–I) Spatial maps and UMAP from the integrated dual-species dataset combining 350 *Medicago* genes and 50 rhizobial genes. Twenty-one clusters were identified; nodule inner cortex (upper; NIC_U, cluster 0), nodule inner cortex (lower; NIC_L, cluster 1), root inner cortex (RIC, cluster 2), nodule middle cortex (lower; NMC_L, cluster 3), uninfected cells in the nitrogen-fixation zone (UninNF, cluster 4), infection zone (IFZ, cluster 5), nodule outer cortex and epidermis (NOC_Epi, cluster 6), infected cells in the nitrogen-fixation zone (InNF, cluster 7), root outer cortex (ROC, cluster 8), nodule uninfected cells and epidermis (NUnin_Epi, cluster 9), lower nodule outer cortex and epidermis (NOC_L_Epi, cluster 10), infected cells in the infection zone (InIFZ, cluster 11), uninfected cells in the infection zone (UninIFZ, cluster 12), pre-infection zone (PreIFZ, cluster 13), upper nodule outer cortex and epidermis (NOC_U_Epi, cluster 14), nodule vasculature endodermis (NV_Endo, cluster 15), nodule meristem and upper nodule inner cortex (NM_NIC_U, cluster 16), uninfected cells in the interzone (UninIZ, cluster 17), infected cells in the interzone (InIZ, cluster 18), infected cells spanning the interzone and nitrogen-fixation zone (InIZ_NF, cluster 19), and senescence zone (SenZ, cluster 20). (J–M) Spatial expression of rhizobial genes across mature nodules. *nifH* is enriched in the interzone, whereas *nifA* is enriched in the nitrogen-fixation zone and reduced in senescence-associated regions. See also Figure S5-7 and Table S1.

Clustering of the 3D spatial transcriptomic dataset resolved major cell types spanning nodule meristematic, cortical, vascular, infection and nitrogen-fixation zones, and late-stage uninfected populations (Figure 2A-B). Unlike 14-DPI nodules, late-stage 49-DPI nodules exhibited pronounced spatial stratification along the longitudinal axis, including distinct cortical and vascular-associated populations that were more readily resolved through 3D reconstruction (Figure 2A-B, Figure S3). This layered organization suggests progressive functional specialization across the apex-to-basal axis and highlights the importance of 3D profiling for resolving structures that are incompletely represented in single two-dimensional sections.

The 3D atlas resolved transcriptionally distinct infected and uninfected populations across infection, interzone, and nitrogen-fixation regions in 49-DPI nodules (Figure 2C–F). Infected cells within the infection-zone (cluster 16) and infected interzone cells (cluster 17) expressed *Early Nodulin 12* (*MtENOD12*), *LONELY GUY 1* (*MtLOG1*), and *Nodule-specific Glycine-rich Protein 1B* (*MtNodGRP1B*), whereas adjacent uninfected cells (cluster 10) expressed *Lectin 4* (*MtLEC4*) (Figure 2C–D; Figure S4).

Two transcriptionally distinct uninfected populations also resolved within the nitrogen-fixation zone that were not resolved at 14 DPI (Figure 2A–F). In addition to infected *MtLb2*-positive cells (cluster 7), an upper uninfected population (cluster 4) expressed *Asparagine synthetase 2* (*MtAS2*) and *Nitrate reductase 2* (*MtNR2*), consistent with active nitrogen assimilation and transport, whereas a second lower and basal population (cluster 11) showed reduced expression of these markers, indicating functional divergence along the longitudinal axis (Figure 2E–F). Uninfected cells associated with the infection and interzone regions (cluster 10) were also transcriptionally distinct from nitrogen-fixation-zone uninfected populations despite limited anatomical separability (Figure 2A–F).

Label transfer from a 14-DPI single-cell atlas to the 49-DPI Xenium dataset showed partial correspondence for nitrogen-fixation-zone populations but limited correspondence for many cortical, vascular, and late-stage populations, highlighting both developmental divergence beyond 14-DPI nodules and changes in transcriptional profiles during nodule maturation that led to limited label transferability (Figure S5). Cell-wall and nuclear-based segmentation produced broadly concordant clusters and markers while differing primarily in regional sensitivity (Figure 2, Figure S5-7). Overall, these analyses demonstrate that mature nodule cell identity is spatially organized, dynamically maintained, and continues to diversify at late stages. However, infected nitrogen-fixation-zone cells remain largely indistinguishable at the host transcriptomic level, motivating dual-species spatial profiling to resolve host–microbe functional states *in situ*.

### Dual-species spatial transcriptomics resolves rhizobial transcriptional states within the nitrogen-fixation zone

To resolve functional heterogeneity within infected cells, we applied Xenium profiling to both the plant host and its bacterial symbiont by incorporating 50 rhizobial targets into dual-species spatial panels (Table S1). The integration of plant and rhizobial transcripts revealed additional transcriptional states within infected tissues that were not detectable using *M. truncatula* targets alone (Figure 2G–I). Rhizobial transcripts contributed to clustering within infected populations, indicating distinct bacterial transcriptional states within morphologically similar host cells (Figure 2G–M). Spatial mapping revealed transitions in bacterial transcriptional states across mature nodules. The nitrogenase structural gene *nifH* was enriched in the interzone, associated with the onset of bacteroid differentiation, whereas the nitrogen-fixation regulator *nifA* was enriched in mature nitrogen-fixation regions (Figure 2J–K), marking the fully established anaerobic state for nitrogen-fixation in mature infected cells.

Dual-species profiling also revealed heterogeneity among nodules at 49 DPI. While some nodules exhibited prominent interzone activity with strong *nifH* expression, others contained extended regions with reduced *nifA* expression, corresponding to a senescence zone, suggesting declining rhizobial activity (Figure 2L–M). These patterns are consistent with observations from STPT imaging (Figure S5C-D, Video S1). Co-expression analysis identified modules linking host and bacterial genes. One module showed spatial co-expression between *MtLb2* and rhizobial genes, including *gltA*, *nifA*, and *fixK1*, consistent with coordinated regulation of oxygen buffering, bacterial respiration, and nitrogen fixation. The same module also contained genes associated with dicarboxylate utilization and central carbon metabolism, indicating coordinated host–microbe transcriptional programs within the nitrogen-fixation zone.

Collectively, these results show that dual-species spatial transcriptomics resolves bacterial transcriptional states within host cells and reveals coordinated host–microbe gene expression that is inaccessible to host-only approaches.

### A time-resolved spatial transcriptomic atlas reveals shared and organ-specific cell identities across nodule and lateral root development

Nodule development builds upon a pre-existing lateral root developmental program but progressively diverges to establish distinct organ identities and morphology. To define the shared and organ-specific developmental trajectories underlying this transition, we constructed wild-type lateral-root and nodule datasets across a continuous developmental series and two ecotypes (A17 and R108) (Figure 3, Figure S8-9). We profiled lateral roots across five developmental stages (1-6 days post induction, DPI) using gravity induction and nodules across nine stages spanning the full trajectory of organogenesis (1-49 DPI) using rhizobial spot inoculation (Figure S8-9; Table S1), enabling direct comparison between the two organs. Integrated analyses were performed using nuclear-anchored segmentation to ensure robust cell detection across densely packed early primordium tissues (Figure 3, Figure S8-9).

**Figure 3.**
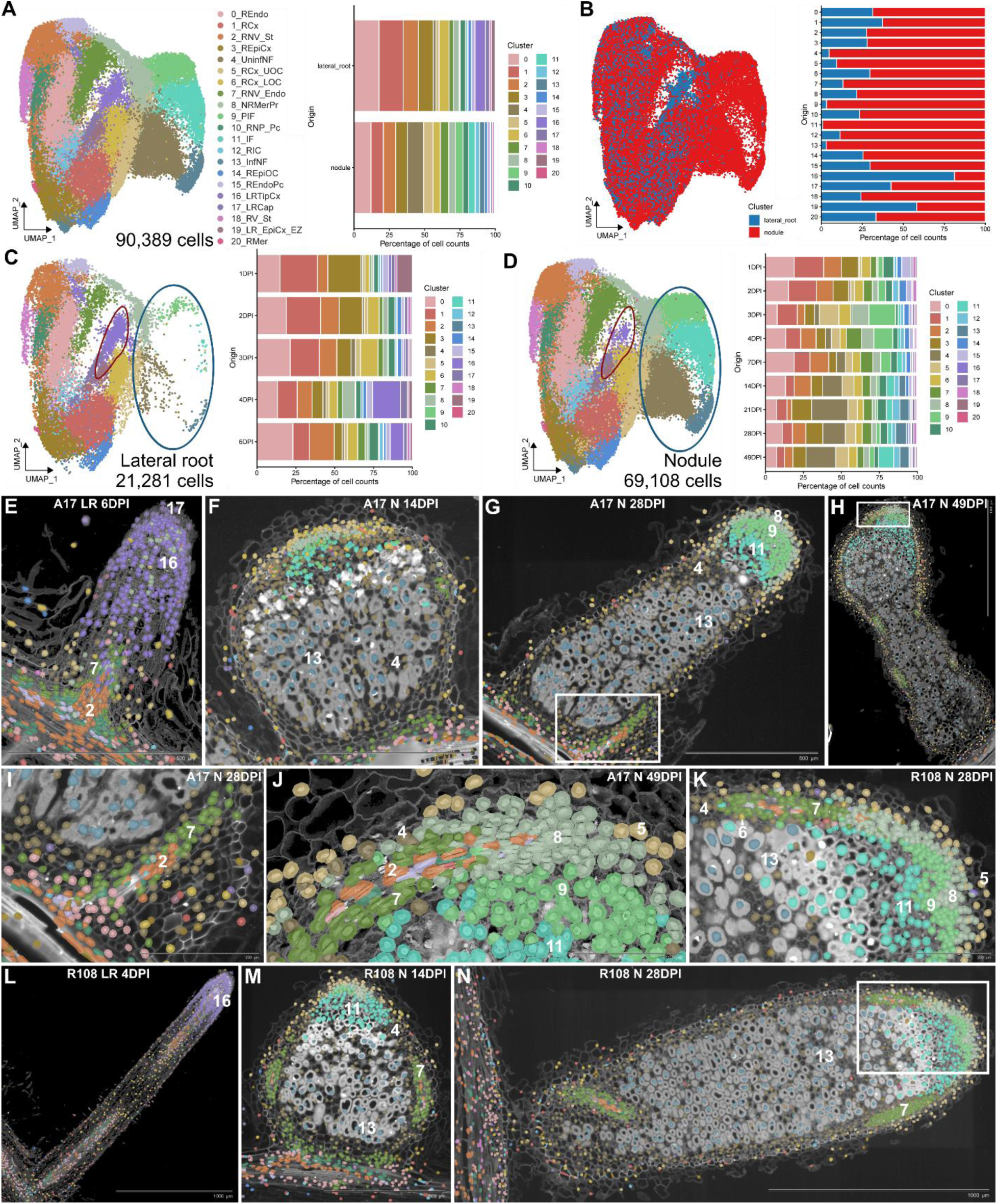
A 4D spatial reference defines shared and organ-specific cell identities across mature lateral roots and nodules. (A) UMAP of the integrated 4D spatial transcriptomic atlas comprising all wild-type lateral root and nodule samples across developmental time and ecotypes (A17 and R108), generated using the 405-gene panel. Twenty-one spatially defined populations were identified; cluster 0, Root endodermis (REndo); cluster 1, Root cortex (RCx); cluster 2, Root + nodule vasculature (stele) (RNV_St); cluster 3, Root epidermis + cortex (REpiCx); cluster 4, Uninfected cells in the nitrogen-fixation zone (UninfNF); cluster 5, Root cortex + nodule upper outer cortex (RCx_UOC); cluster 6, Root cortex + nodule lower outer cortex (RCx_LOC); cluster 7, Root + nodule vasculature (endodermis) (RNV_Endo); cluster 8, Nodule + root meristem/primordium (NRMerPr); cluster 9, Pre-infection zone (PIF); cluster 10, Root + nodule pericycle (RNP_Pc); cluster 11, Infection zone (IF); cluster 12, Root inner cortex (RIC); cluster 13, Infected cells in the nitrogen-fixation zone (InfNF); cluster 14, Root epidermis + outer cortex (REpiOC); cluster 15, Root endodermis + pericycle (REndoPc); cluster 16, Lateral root tip cortex (LRTipCx); cluster 17, Lateral root cap (LRCap); cluster 18, Root vasculature (stele) (RV_St); cluster 19, Lateral root epidermis + cortex (elongation zone) (LR_EpiCx_EZ); cluster 20, Root meristem (RMer). Right, bar plot showing cluster composition by organ. (B) UMAP colored by tissue type, highlighting organ-enriched clusters. (C–D) UMAPs of lateral-root cells (D) and nodule cells (E), with bar plots showing cell-type composition across developmental stages. Nodule-enriched clusters are indicated by blue outlines and lateral-root–enriched clusters by red outlines. (E–H) Spatial maps of A17 tissues colored by cell-type identity: lateral root at 6 DPI (E), and nodules at 14, 28, and 49 DPI (F–H). (I–K) Magnified mature nodule regions showing basal vasculature (I), meristem-associated vasculature (J) and the meristem-to-infection-zone transition (K). (L–N) Spatial maps of R108 tissues colored by the same cell-type identities: lateral root at 4 DPI (L), and nodules at 14 and 28 DPI (M–N). See also Figure S8-9 and Table S1.

Joint embedding of all wild-type lateral root and nodule samples generated a unified UMAP comprising 90,389 filtered cells and 21 spatially defined clusters spanning shared and organ-specific identities (Figure 3A; Table S1). While many clusters contained cells from both organs (Figure 3B), infection-related populations (cluster 9, 11), and nitrogen-fixation zone cells (cluster 4, 13) were exclusively nodule-derived. In contrast, lateral root tip cortex (cluster 16), lateral root cap (cluster 17), and elongation-zone epidermis and cortex (cluster 19) were restricted to lateral roots (Figure 3A–D). Temporal analysis revealed contrasting developmental dynamics (Figure 3C–D): lateral root identities were established rapidly (4–6 days), whereas nodules exhibited progressive, stage-dependent remodeling, marked by sequential emergence of pre-infection, infection, and nitrogen-fixation–zone populations. Despite these differences, the majority of cell types show overlapping transcriptomic profiles between lateral roots and nodules, highlighting the commonality of these developmental structures, with specific divergence driven by unique transcriptional programs in defined cell populations (Figure 3A–D).

Spatial mapping resolved distinct lateral-root, infection-zone, and nitrogen-fixation-zone populations and revealed how shared and organ-specific cell identities are organized during organ development (Figure 3E-H). High-resolution views further revealed shared transcriptional features between lateral root and nodule vasculature, with stele and endodermal populations near the nodule base (cluster 2, 7) showing identities closely related to primary root tissues (Figure 3E, I). Vascular-associated populations were also consistently detected near the nodule apex adjacent to a shared root–nodule meristem/primordium population (cluster 8) (Figure 3J), which was observed at both lateral root tips and nodule apices across developmental stages and ecotypes (Figure 3E–N). In mature nodules (14-49 DPI), a meristematic population (cluster 8) was consistently positioned adjacent to pre-infection and infection-associated populations (cluster 9, 11), vascular-associated endodermal cells (cluster 7), and outer cortical and uninfected nitrogen-fixation–zone populations (cluster 5, 6, 4) (Figure 3J-K). The spatial juxtaposition of these cell types supports an organized architecture centered on a single meristematic domain from which multiple lineage programs emerge.

To determine when lineage relationships observed in mature nodules first emerge, we analyzed early developmental stages of lateral root and nodule primordia to compare their initial spatial and transcriptional organization (Figure 4). Early lateral root development (1–4 DPI) exhibited a comparatively homogeneous organization (Figure 4A–E), with primordia dominated by a shared meristematic population (cluster 8) together with vascular-associated cell types (cluster 7, 10, 2) (Figure 4A–E). By 6 DPI, lateral roots had differentiated toward lateral root–specific cortex and tip identities (cluster 16) while maintaining prominent vascular-associated populations. Overall, lateral root development followed a relatively linear trajectory, with early establishment of core identities and limited diversification.

**Figure 4.**
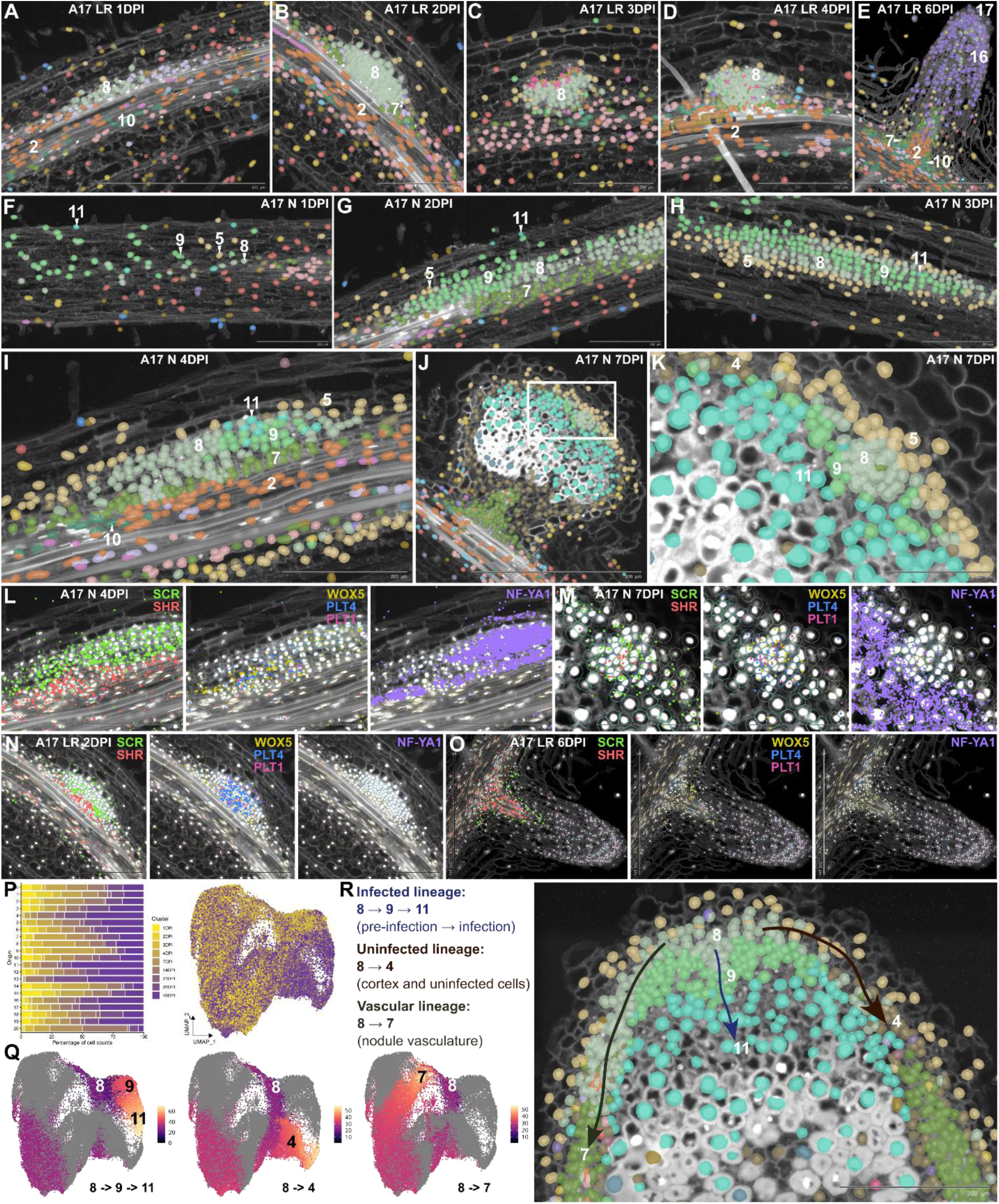
Early spatial diversification in nodule primordium reveals multiple developmental programs underlying nodule organogenesis. (A–E) Spatial maps of A17 lateral roots at 1, 2, 3, 4, and 6 DPI, colored by the cell-type identities defined in Figure 3. (F–J) Spatial maps of A17 nodules at 1, 2, 3, 4, and 7 DPI, showing emergence of meristematic (8), cortical (4), pre-infection (9), and infection-zone (11) populations. (K) Magnified 7-DPI nodule tip showing spatial organization of meristematic tissue (8), pre-infection zone (9), infection zone (11) and surrounding cortical populations. (L–O) Spatial expression of representative developmental regulators during early nodule and lateral-root development. *SCR* and *SHR* mark endodermal, cortical, and vascular-associated domains; *WOX5*, *PLT1*, and *PLT4* mark meristematic populations; *NF-YA1* is enriched in nodule pre-infection and infection-associated tissues but absent from lateral-root primordia. (P) UMAP colored by true developmental time (1–49 DPI) with corresponding stage-resolved cell-type composition. (Q) Slingshot pseudo-time trajectories from the shared primordium/meristem population toward vascular, cortical/uninfected, and infection-associated lineages. Cells are colored by pseudo-time. (R) Conceptual model summarizing three major developmental programs emerging from the shared nodule meristem: infected, uninfected, and vascular lineages. See also Figure S10.

In contrast, early nodule primordia exhibited substantially greater heterogeneity from the earliest stages analyzed (1–4 DPI) (Figure 4F–I). Multiple transcriptionally distinct populations emerged around the shared primordium/meristem (cluster 8), including pre-infection, infection-associated (cluster 9, 11), cortical (cluster 5), and vascular-related cells (cluster 7, 10) that foreshadow mature nodule organization. Although morphologically indistinguishable at these stages, these cell populations already display distinct transcriptional identities, indicating that lineage diversification precedes overt anatomical zonation. By 7 DPI, these domains are resolved into an organized meristematic structure surrounded by infection-associated, cortical, and vascular-associated tissues (Figure 4J-K). Together, these observations suggest that, unlike lateral roots, nodules initiate multiple developmental programs in parallel during early primordium formation.

Marker gene expression further supported these differences (Figure 4L–O). *SCR* and *SHR* expression highlighted endodermal and pericycle domains in lateral root primordia, but in nodules, *SCR* extended into cortical and nodule primordia-associated regions (Figure 4L, N), consistent with previous reports of expanded *SCR* expression in the cortex during nodulation^21^. Meristem-associated regulators (*WOX5*, *PLT1* and *PLT4*) displayed asymmetric organization in nodule primordia compared with the compact centralized domains observed in lateral root primordia (Figure 4L, N, Figure S10), reflecting the peripheral positioning of developing nodule vasculature rather than a single central axis. *NF-YA1* was absent from lateral root primordia but enriched in nodule primordia, localizing to cells adjacent to the nodule primordia and overlapping with emerging pre-infection and infection-associated populations (Figure 4L-O). These patterns indicate that nodule primordia comprise coexisting lateral root-like and nodule-specific transcriptional programs in different cell populations.

To relate spatial organization to developmental progression, we integrated true developmental time with Slingshot pseudo-time inference (Figure 4P-Q, Figure S9D). In lateral roots, trajectories from the primordium/meristem cells (cluster 8) primarily resolved toward lateral root tip (cluster 16) and vascular endodermal identities (cluster 7). In nodules, primordium/meristem cells were similarly connected to vascular endodermal identities (cluster 7), indicating that nodule vascular development likely adopts a root-derived developmental program. Nodule primordia generated additional trajectories toward cortical and uninfected nitrogen-fixation–zone cells (cluster 4), infection-zone cells (cluster 11), and infected nitrogen-fixation–zone cells (cluster 13) that were not observed in lateral roots. Integrating spatial, temporal, and trajectory analyses supports a model in which a common primordium/meristem population gives rise to three coordinated developmental programs: vascular, non-symbiotic cortical, and symbiotic infection-associated programs (Figure 4R). Together, these results show that nodule cell-type diversification is established at the earliest stages of primordium formation, with multiple concurrent developmental cell lineages to support symbiotic function, as well as a conserved program with lateral roots that leads to vascular development. Importantly, the cell lineage structure inferred from mature nodules also reflects early and persistent patterning decisions, providing a conceptual framework for interpreting developmental trajectories.

### The role of nodulation regulators in the establishment of nodule development and cell-type specification

The comparison between nodules and lateral roots demonstrates unique cellular developmental trajectories that give rise to the key functions of nodules: cells that can accommodate nitrogen-fixing bacteria and uninfected cells that appear to function in nitrogen assimilation. Already well-established genetic regulators have been shown essential for nodule function, *NIN, CRE1, NF-YA1, LSH1/LSH2* and *NOOT1/NOOT2*, as well as regulators that show overlapping functions in lateral roots and nodules, *LBD16* and *STY*. To better understand the function of these regulators we first mapped their expression patterns at early developmental stages (4 and 7 DPI; Figure 5A–B; Figure S11).

**Figure 5.**
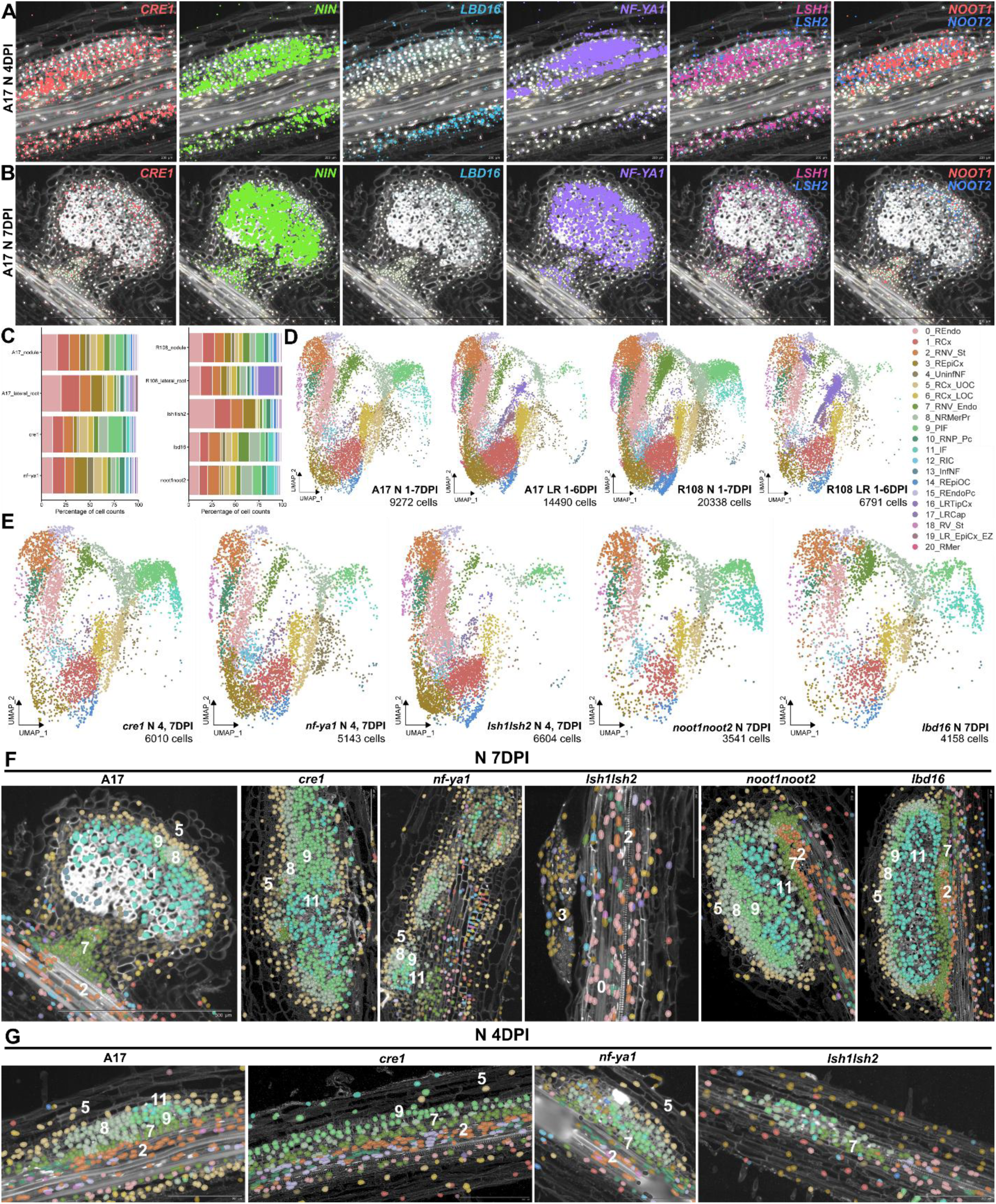
Perturbation of key nodulation regulators and cytokinin signaling alters early nodule cell-type emergence and spatial organization. (A–B) Spatial distribution of *CRE1*, *NIN*, *LBD16*, *NF-YA1*, *LSH1/LSH2* and *NOOT1/NOOT2* in representative A17 nodules at 4 DPI (A) and 7 DPI (B). (C) Cluster-resolved cell-type composition across wild-type and mutant tissues. Colors correspond to the spatially defined clusters shown in Figure 3. (D–E) Integrated UMAPs and cluster composition of wild-type (D) and mutant (E) datasets across early developmental stages.\ (F–G) Spatial maps of wild-type and mutant nodules colored by cluster identity. (F) A17, *cre1, nf-ya1, lsh1/lsh2, noot1/noot2,* and *lbd16* at 7 DPI. (G) A17, *cre1, nf-ya1,* and *lsh1/lsh2* at 4 DPI. See also Figure S11-12.

At 4 DPI, consistent with the established role of cytokinin perception in early nodule initiation^17,18^, *CRE1* and *NIN* were co-enriched in inner cortex, endodermis, and pericycle regions of the primordium (Figure 5A, Figure S11), together with *NF-YA1*, *LSH1/LSH2*, and *NOOT1/NOOT2*, defining a central regulatory axis that demarcates nodule identity from surrounding root tissues. *NF-YA1* was enriched in pre-infection and infection-associated cells but excluded from vascular precursors (Figure 5A), reinforcing the early segregation of vascular and infection lineages (Figure 4). In contrast, *LBD16* was restricted to outer cortical layers (Figure 5A), suggesting a lateral-root-like cortical program that is partially redeployed during nodule initiation but remains spatially segregated from the cytokinin–*NIN* axis.

By 7 DPI, these regulatory domains become progressively refined (Figure 5B, Figure S11). *CRE1* remained broadly distributed but showed relative enrichment near the meristem and vascular-associated regions. *NIN* and *NF-YA1* were concentrated within the pre-infection and infection zones, aligning with their established roles in coordinating infection and symbiotic differentiation, with *NF-YA1* showing its strongest enrichment in the pre-infection and infection zones (Figure 5B), consistent with previous observations that *NF-YA1* expression is confined to infection-associated tissues as nodules mature^14^. Conversely, *LBD16* expression was reduced and confined to a narrow apical domain adjacent to developing vasculature, supporting its primary function during early primordium establishment rather than later zonation (Figure 5B). Surprisingly, the nodule-identity regulators *LSH1/LSH2* and *NOOT1/NOOT2* were enriched in meristematic tissues and peripheral uninfected cells surrounding developing infection and nitrogen-fixation zones (Figure 5B, Figure S11), suggesting that maintenance of nodule identity requires coordinated regulation from peripheral domains rather than infected cells or meristem alone. Collectively, these data reveal a modular architecture comprising a central cytokinin–*NIN* axis, an infection-associated module, and a peripheral identity-maintenance module that collectively define early nodule patterning.

To test how these modules control organogenesis, we integrated their loss-of-function mutants into the spatial atlas, focusing on *cre1*^18^ and *nf-ya1*^37,38^ (A17 background), and *lsh1/lsh2*^23^, *noot1/noot2*^39^, and *lbd16*^10^ (R108 background) (Figure 5C–G). We excluded *nin* from these analyses, since we already know that loss-of-function *nin* mutants abolish almost the entirety of nodulation-associated gene expression and lack nodule initiation^40^.

*cre1* and *nf-ya1* retained most early nodule-associated clusters but differed in their relative abundance. *cre1* nodules showed a proportional increase in pre-infection and early infection-related populations (cluster 9, 11) (Figure 5C-G), consistent with previous reports and our observation of hyperinfection phenotypes (Figure 5F-G; Figure S12A) at early stages in *cre1* mutants^18^. In contrast, *nf-ya1* nodules displayed reduced infection-related clusters and primordium/meristem-associated cells (cluster 8), in agreement with observations that *nf-ya1* mutants exhibit restricted infection (Figure 5F-G; Figure S12A) and reduced or absent meristematic activity ^14,37^.

At later timepoints, *cre1* nodules became progressively disorganized (Figure 6A–D). Although early stages showed enhanced infection, mature *cre1* nodules displayed substantial variation in size and some failed to maintain coherent infection zones (Figure 6D). Some *cre1* nodules were unable to sustain a persistent meristematic domain superficially resembling a determinate nodule with a centralized nitrogen-fixation zone but limited zonation characteristic of wild-type *M. truncatula* nodules. While integrated analyses revealed enrichment of lateral-root–associated populations, spatial mapping demonstrated that these cells originated from adjacent lateral roots rather than the nodule itself (Figure 6A–D), indicating that *cre1* does not reprogram nodule tissue toward lateral-root fate. Instead, uninfected cortical and vascular-associated populations expanded, whereas symbiotic lineages failed to stabilize, suggesting that *CRE1*-dependent cytokinin signaling is required to reinforce nodule differentiation and maintain the nodule meristem.

**Figure 6.**
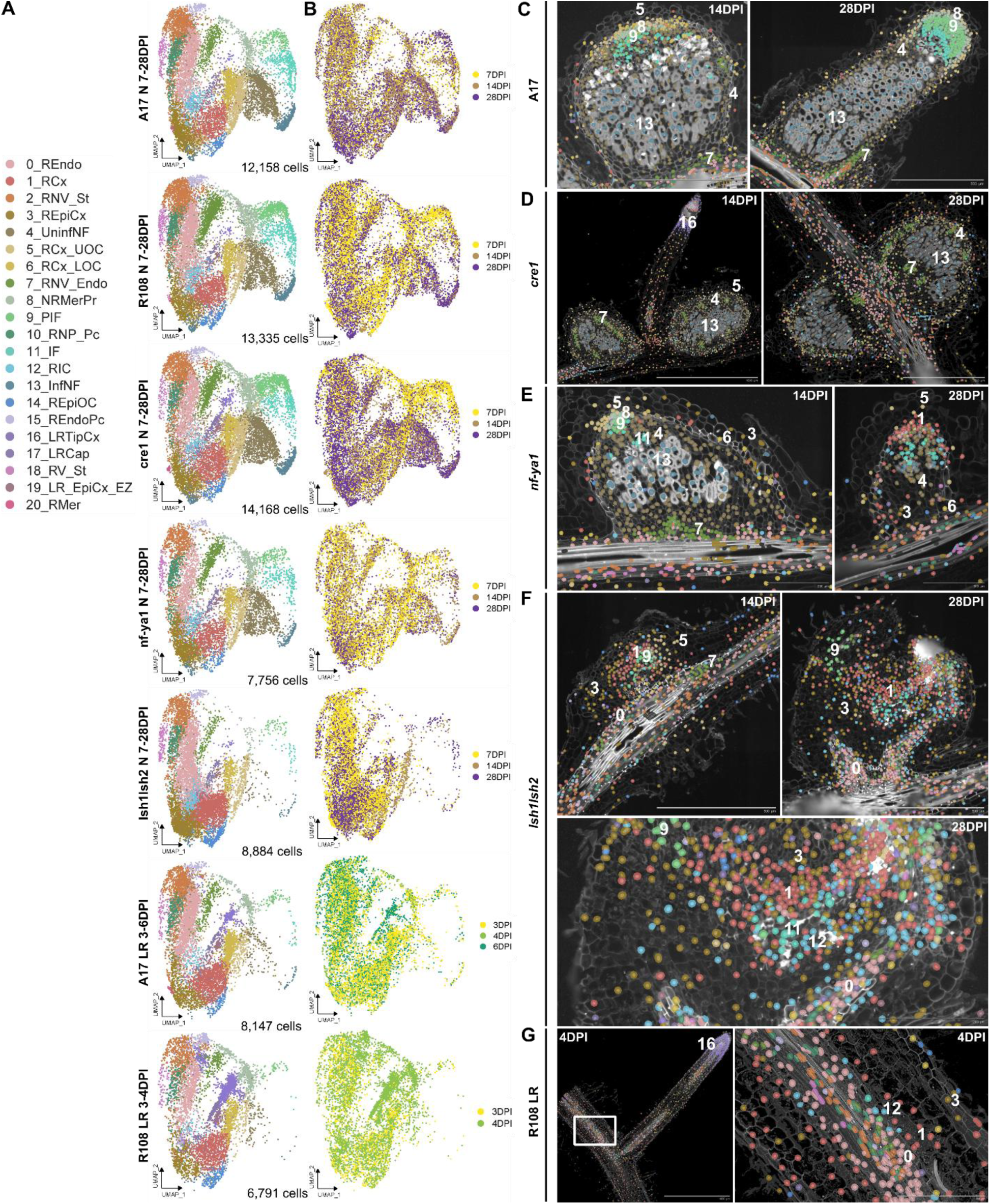
Later-stage mutant profiling reveals collapse toward primary-root-associated states. (A) Integrated UMAPs of wild-type A17/R108 nodules, *cre1*, *nf-ya1* and *lsh1/lsh2* nodules, and wild-type A17/R108 lateral roots across developmental time, colored by Figure 3 cell-type identities. (B) Same UMAPs are colored by developmental time. (C–F) Spatial maps of mutant nodules at 14 and 28 DPI: (C) A17, (D) *cre1*, (E) *nf-ya1*, and (F) *lsh1/lsh2*. Bottom, magnified view of the 28-DPI *lsh1/lsh2* nodule showing central tissue organization. (G) Spatial maps of R108 lateral and primary roots at 4 DPI for comparison with *lsh1/lsh2* nodules. See also Figure S12.

Similarly, *nf-ya1* nodules exhibited restricted infection, loss of meristematic activity, and overall reduced nodule size (Figure 6A-C, E). During maturation, infection-associated and meristematic populations (cluster 9, 11, 8) progressively declined (Figure 6A-C, E), whereas uninfected nitrogen-fixation–zone identities (cluster 4) and primary-root–associated cortical populations (cluster 3, 1, 6) expanded within the nodule structure. By 28 DPI, *nf-ya1* nodules were smaller and frequently lacked a coherent meristematic domain, with severely reduced infection and nitrogen-fixation zones. True-time embeddings and pseudo-time analyses showed developmental stalling, with few cells progressing to mature symbiotic states (Figure 6B, Figure S12). These findings indicate that *NF-YA1* is required for both infection progression and stabilization of nodule identity during maturation. In its absence, cortical cells partially retain a primary root–like transcriptional state. Thus, both *cre1* and *nf-ya1* nodules exhibited attenuation of the meristematic program and a reduction in cell populations associated with mature nodule function (Figure 6A–E), although all major nodule cell types remained detectable.

The most severe phenotype was observed in *lsh1/lsh2* mutants, which showed near-complete loss of nodule-specific populations, including both infected and uninfected nitrogen-fixation–zone lineages (cluster 13, 4), pre-infection and infection populations, and the primordium/meristem population (Figure 5C-G), the cell-types that define nodule functionality. This loss was accompanied by an increased contribution of transcriptional states associated with root tissues (clusters 3) (Figure 5F), consistent with previous reports that *lsh1/lsh2* mutants fail to maintain nodule identity and display abnormal, root-like morphology^23^. Notably, despite this collapse of nodule identity, *lsh1/lsh2* nodules did not acquire lateral-root–specific cortex identities (cluster 16) (Figure 5C-E), rather *lsh1/lsh2* nodules were dominated by primary-root–like transcriptional identities (cluster 3) throughout the nodule primordium-like structure (Figure 5F). At late time-points, the structures that form on *lsh1/lsh2* mutants lack any of the developmental structure observed in wild-type, with a collapse in the organization and presence of nodule-associated cell types and the incursion of cell-types normally restricted to the primary root (Fig. 6A-B, F-G). This indicates that loss of nodule identity reflects a failure to complete or maintain the transition from primary root identity to nodule identity, rather than reprogramming toward lateral root fate.

In contrast, *noot1/noot2* and *lbd16* mutants showed very subtle phenotypes: most early nodule cell types were retained, but selective perturbations in specific clusters led to modest shifts in cell-type proportions rather than wholesale collapse of nodule identity (Figure 5C-F; Figure S12A).

Together, these analyses differentiate regulator functionality: nodule meristem maintenance and expansion (CRE1 and *NF-YA1),* initiation of nodule identity and functionality (*LSH1/2*), and the proliferation of tissues/cells within the emerging nodules (*NOOT1/2* and *LBD16*).

### Hormone signaling during nodule organogenesis

The hormones auxin, cytokinin and gibberellins (GAs) are emerging as key regulators of nodule development^1,2,23,24^ and therefore we included components that control and respond to these hormones within our selected gene probes in wild-type and mutant nodules (Figure 7; Figure S13). In wild-type nodules, meristem marker genes and hormone signaling genes form a coherent, progressively refined spatial expression architecture (Figure 7; Figure S13). At 4 DPI, *WOX5* broadly marks the primordium, while *PLT1* becomes restricted to vascular-associated domains (Figure S13A). By 14 DPI, both converge within a compact meristem (Figure 7A; Figure S13A-B). Auxin-associated genes exhibit complementary specificity: *STY4* broadly marks the early primordium and remains enriched in meristematic and vascular tissues, whereas *YUC8* becomes increasingly restricted to meristem- and vascular-associated domains (Figure 7A; Figure S13A-B). The Cytokinin activation gene *LOG1* is initially broad but becomes concentrated near the meristem and vasculature (Figure 7A; Figure S13A-B), while cytokinin response (*RR4*) sharpens at the interzone (Figure 7A; Figure S13A-B), consistent with its role in coordinating the transition from infection to fixation. GA biosynthesis and response genes establish a longitudinal axis aligned with zonation, with *GA3ox1* and *GA3ox2* enriched in infection zones and *GASA8* and *GASA22* marking non-infected fixation-associated domains (Figure 7A; Figure S13A-B). Together, these patterns define a spatially integrated meristem–hormone regulatory framework, with auxin domination in the meristematic zone, cytokinin domination in the infection zone and interzone, and at least as it pertains to uninfected cells GA dominating in the fixation zone.

**Figure 7.**
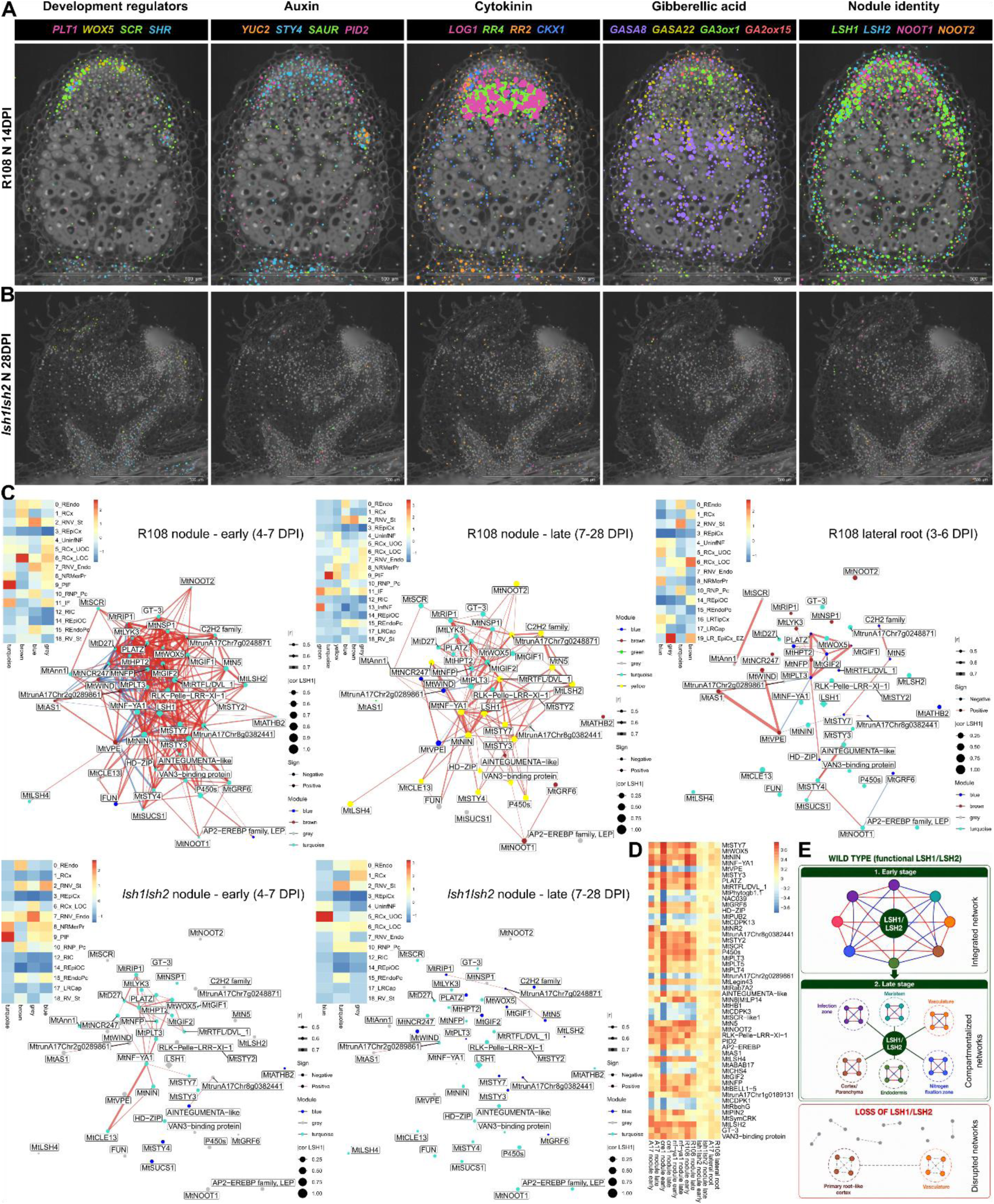
Collapse of coordinated hormone–meristem programs and rewiring of gene co-expression architecture in *lsh1/lsh2* nodules. (A–B) Spatial visualization of meristem and hormone-related transcripts in wild-type R108 nodules at 14 DPI (A) and *lsh1/lsh2* nodules at 28 DPI (B). Genes are grouped by functional category: transcription factors that are developmental regulators (*PLT1*, *WOX5, SCR1*, *SHR1*), auxin biosynthesis and regulation (*YUC8*, *STY4*) and response (*Small Auxin Upregulated DNA, SAUR-like*) and transport (*PINOID-like*, *PID2*), cytokinin biosynthesis and response (*LOG1*, *RR4, RR2, CKX1*), gibberellin metabolism and response (*GASA8*, *GASA22*, *GA3ox2*, *GA2ox15*), and nodule identity regulators (*LSH1, LSH2, NOOT1, NOOT2*). (C) *LSH1*-centered signed gene co-expression networks showing developmental and genotype-dependent changes in regulatory architecture. Networks were constructed using genes selected from the wild-type R108 nodule reference network and displayed using a fixed network layout to enable direct comparison across conditions. Networks are shown for early- and late-stage R108 nodules, R108 lateral roots, and early- and late-stage *lsh1/lsh2* nodules. Node size reflects the absolute correlation with *LSH1*, and node color indicates hdWGCNA module assignment. Red and blue edges indicate positive and negative gene-gene correlations, respectively, with edge thickness proportional to the absolute Pearson correlation coefficient. Companion module eigengene (ME) heatmaps show the average expression of each co-expression module across cell clusters in the corresponding condition. Columns represent hdWGCNA modules and rows represent cell clusters as defined in Figure 3A. (D) Heatmaps showing Pearson correlation coefficients between *LSH1* and the top *LSH1*-associated genes across developmental stages, genotypes, and organs. Gene sets were selected from the wild-type R108 nodule reference network and compared across wild-type and mutant nodules (R108, *lsh1/lsh2*, *nf-ya1*, and *cre1*) as well as lateral roots. (E) A model of *LSH1/2*-centered network dynamics during nodule development. Early nodules exhibit a highly integrated regulatory network that becomes progressively compartmentalized into cell-type-specific modules during maturation. *LSH1/2* remains connected across these modules, consistent with a role in coordinating diverse developmental programs. In *lsh1/lsh2* mutants, network connectivity is reduced at early stages and largely collapses at later stages, indicating a requirement for *LSH1/2* in establishing and maintaining coordinated cell-type-specific regulatory networks. See also Figure S13.

Mutations in *cre1* show an absence of meristematic tissues in mature nodules, as previously described, and thus lack the markers *WOX5, PLT1* and meristem-associated *STY4* and *GA2ox15* expression, although auxin-associated vascular expression patterns remained (Figure S13C). GA-associated genes lost their normal zonation, indicating that *CRE1*-dependent cytokinin perception is required to stabilize meristem identity and likely coordinate downstream hormone patterning. In contrast, *nf-ya1* nodules retained the overall hormone architecture but in an attenuated form. A small meristematic domain persisted, marked by confined *WOX5* and *PLT1* expressions at the nodule apex, while auxin-associated genes remained enriched in the vasculature and *RR4* was restricted to a narrow domain near the fixation zone (Figure S13D). In contrast, *lsh1/lsh2* mutants show a complete collapse in the hormone program (Figure 7A-B), with no coherent pattern in the hormonal landscape of *lsh1/lsh2* nodules. Strikingly, although hormone-related transcripts remained detectable in subsets of cells, their expression was largely confined to vascular and root-like tissues and failed to form the coherent spatial domains observed in wild-type nodules (Figure 7A-B).

Collectively, these data show that nodule organogenesis is associated with precise spatial coupling between meristem identity and hormone signaling. Disruption of cytokinin perception or *LSH1/LSH2* progressively uncouple these programs, leading to meristem loss, the collapse of hormone gradients and the collapse of nodule identity and structure.

### Dynamic network analysis reveals *LSH1/LSH2*-dependent coordination of developmental regulatory programs during nodule organogenesis

From the work described above it is apparent that *LSH1/LSH2* are the most critical components for nodule identity and functionality. To investigate how *LSH1* contributes to nodule development, we constructed *LSH1*-centered signed gene co-expression networks and compared network architecture across developmental stages, genotypes, and organs (Figure 7C–D, Figure S12). By separating early (4–7 DPI) and late (7–28 DPI) developmental stages, this framework enabled direct comparison of network rewiring during nodule maturation.

In wild-type R108 nodules, early developmental stages were characterized by a highly interconnected co-expression network linking regulators of nodulation (*LSHs*, *NOOTs*, *NIN*, *NF-YA1*), hormone signaling (*STY*s), meristem maintenance (*WOX5*, *PLT*s), and developmental patterning (*SCR*) (Figure 7C-D). Most of these genes were co-expressed within the same module, reflecting an integrated regulatory program operating during early nodule organogenesis. As nodules matured, this integrated network became compartmentalized into distinct co-expression modules associated with specific cell types and developmental domains (Figure 7C). Despite this increasing specialization, *LSH1* remained connected across multiple modules, suggesting a role in coordinating diverse developmental programs throughout the nodule (Figure 7C). This interpretation is consistent with the broad peripheral expression pattern of *LSH1*, which spans multiple nodule domains, including the meristem, infection zone, cortex, and uninfected cells of the nitrogen-fixation zone (Figure 7A).

The network architecture was substantially disrupted in *lsh1/lsh2* nodules. Many co-expression relationships present in wild-type nodules were already weakened or absent during early development and were largely lost by late stages in the mutant (Figure 7C-D). However, connections involving *NIN*, *NF-YA*, *WOX5*, and *PLT* genes remained detectable in the early-stage mutant network, suggesting that these pathways may act upstream of, in parallel with, or partially independent of *LSH1/LSH2* (Figure 7C). At the same time, a large proportion of the wild-type developmental network was already disrupted at these early stages, indicating that many downstream developmental programs depend on *LSH1/LSH2* activity from the onset of nodule organogenesis. The extensive *LSH1*-centered connectivity observed in wild-type nodules was absent from both *lsh1/lsh2* nodules and lateral roots, although *PLT*- and auxin-associated subnetworks remained present. Importantly, the *lsh1/lsh2* mutant network did not resemble the lateral root network (Figure 7C), indicating that loss of *LSH1/LSH2* does not result in reversion to a lateral root developmental state but instead disrupts the establishment of nodule-specific regulatory programs.

Analysis of A17 nodules and nodulation mutants further supported this interpretation. Although *nf-ya1* and *cre1* mutants exhibited alterations in *LSH1*-associated network architecture, both retained substantially more of the wild-type connectivity than *lsh1/lsh2* nodules. The *nf-ya1* mutant showed partial disruption of the specific *NF-YA1* connected part of network relationships, while preserving much of the broader architecture, whereas *cre1* displayed even more subtle network rewiring, with some connections weakened and others strengthened, suggesting cytokinin signaling may function more as a broader regulatory feedback system. These observations are consistent with the comparatively milder developmental phenotypes of these mutants and support a central role for *LSH1/LSH2* in maintaining coordinated nodule regulatory programs.

Together, these analyses support a model in which early nodule development is driven by an integrated regulatory network that progressively resolves into specialized cell-type-specific programs during maturation (Figure 7E). *LSH1/LSH2* appears to function as a network coordinator linking these developmental modules across multiple nodule domains. In the absence of *LSH1/LSH2*, network organization progressively collapses, resulting in the loss of coordinated cell-type-specific regulatory programs, preventing establishment of the transcriptional architecture required for mature nodule identity. More broadly, these findings highlight how spatial transcriptomics can uncover concurrent regulatory programs operating across distinct developmental domains and reveal the higher-order network architecture underlying organogenesis that is largely inaccessible to bulk transcriptomic analyses. The maintenance of these spatial co-expression relationships appears to be a critical component of nodule identity, developmental patterning, and organ function.

## DISCUSSION

Nodules are unique organs in the plant kingdom that create cells uniquely able to harbor nitrogen-fixing bacteria and provide an environment that supports the action of bacterial nitrogenase^41^. Decades of research has identified genetic regulators controlling nodule initiation and development^2,42^ and revealed shared developmental programs between nodules and lateral roots^1^. However, it remains an open question how nodule identity is established, coordinated, and maintained, and how an underpinning lateral root program is modified to generate the specialized cell types and functions of nodules. By integrating spatial organization with developmental time, our 4D spatial transcriptomic atlas reveals that lateral-root-associated programs are retained within discrete cell types, particularly those associated with vascular development. Nodule-specific functions appear to evolve not through a linear transition from lateral-root to nodule identity, but through the addition of specialized cellular programs that develop and are maintained in parallel. These findings imply that the development and evolution of nodulation required the stacking of developmental trajectories, generating greater cellular complexity than is observed in lateral roots. Strikingly, loss of nodule identity is not a simple switch to a lateral-root fate. Instead, identity-deficient nodules adopt a chimeric organ transcriptional state dominated by primary-root–associated programs, accompanied by the collapse of nodule-specific regulatory modularity rather than linear fate conversion. These findings challenge the prevailing assumption that nodules behave as modified lateral roots whose disruption leads to lateral-root reversion. Rather, our data support a model in which both nodule and lateral-root cell identities are actively maintained developmental states existing in parallel, requiring spatial coordination of regulatory and hormonal programs.

Previous studies have provided important understanding of nodule development through bulk transcriptomic and single-cell studies, identifying major developmental regulators and cell types associated with nodulation at specific developmental stages^23,26,28,43,44^. However, dissociation-based approaches can introduce cellular and transcriptional biases during sample preparation^30,31^, inherently remove spatial context^32,33^, and may underrepresent rare, fragile, or structurally embedded cell populations^31^. Therefore, spatial relationships and lineage progression must be inferred indirectly from marker genes and transcriptional similarity. In plants, cell-type annotation frequently relies on marker genes identified through promoter-reporter studies, *in situ* hybridization, or orthologous markers transferred from model species. However, promoter activity does not necessarily reflect endogenous transcript localization, and gene duplication and functional divergence can alter spatial expression domains even among conserved orthologs^34,35^, complicating marker transfer across species and developmental contexts. By directly measuring transcript localization within intact tissues, spatial transcriptomics provides a spatially anchored framework for cell-type annotation and developmental analysis. Although Xenium profiles a targeted gene set instead of the whole transcriptome, the addition of spatial information at cellular and subcellular resolution improves characterization of cell states, developmental organization, and cell–cell relationships, enabling identification of spatially distinct populations that are difficult to resolve using dissociation-based approaches alone.

Our 4D spatial atlas supports a model in which a shared meristematic domain gives rise to three coordinated developmental programs: symbiotic infection-associated, non-symbiotic cortical/uninfected, and vascular lineages (Figures 1–6). Of these, only the vascular lineage remains transcriptionally aligned with primary-root and lateral-root vasculature, consistent with repurposing of root vascular developmental modules during nodulation (Figures 3 and 4). In contrast, the symbiotic and non-symbiotic lineages are specific to nodule development, whereas lateral roots retain distinct root tip and cortical programs not observed during nodulation. Together, these findings suggest that nodule organogenesis involves the parallel deployment of shared root developmental modules and specialized nodule cell identities.

Spatial transcriptomics provides a new level of developmental resolution and insight that cannot be inferred from morphology or dissociation-based transcriptomic approaches alone. Rather than relying on assumptions about organ identity based on structures observed, spatial mapping directly resolves how developmental regulators influence the emergence, organization, and maintenance of specific cell types. Functional perturbation reveals that nodule identity depends on coordinated regulatory input. Disruption of these regulators produces distinct failure modes, ranging from incomplete stabilization of symbiotic programs in *cre1* to developmental stalling in *nf-ya1* and collapse toward a primary-root-associated state in *lsh1/lsh2*. *CRE1* is revealed to be associated with overall growth and the maintenance of the nodule meristem, although substantial phenotypic variation exists among *cre1* nodules. Likewise, *NF-YA1* function is principally associated with infection-zone development and expansion of the meristematic domain. Previous work showed that *nf-ya1* mutants show severe reduction of the central meristematic domain^45^, and vascular tissues remain largely confined to the nodule base. Because *NF-YA1* expression is enriched in primordium and central meristem-associated domains rather than vascular precursors, the vascular defect in *nf-ya1* likely reflects indirect dependence on an intact central nodule meristematic program.

The most striking phenotype was observed in *lsh1/lsh2*. Although rhizobia-induced structures still form, they fail to establish the coordinated cellular organization and hormone-associated patterning characteristic of wild-type nodules. While previous studies implicated *LSH1/LSH2* in nodule identity, our spatial analyses further reveal the extent of their role in coordinating nodule-specific developmental programs. Strikingly, loss of *LSH1/LSH2* does not result in reversion to a lateral-root identity. Instead, *lsh1/lsh2* structures are dominated by primary-root-associated cell states, indicating that nodule-specific identities are maintained independently of the lateral-root developmental program. These findings support a model in which *LSH1/LSH2* are required for the establishment and maintenance of specialized cellular identities that develop in parallel to, rather than directly from, lateral-root trajectories. Dynamic network analysis further suggests that nodule organogenesis proceeds through a transition from an integrated early developmental program to increasingly compartmentalized cell-type-specific regulatory modules during maturation (Figure 7). Within this framework, *LSH1/LSH2* appear to function as higher-order coordinators that maintain connectivity across otherwise distinct developmental programs. In *lsh1/lsh2* mutants, disruption of this coordinating function is evident from early developmental stages and culminates in the collapse of network architecture at later stages. The persistence of residual connections involving *NIN*, *NF-YA*, *WOX5*, and *PLT* genes suggests that some regulatory pathways may act upstream of, in parallel with, or independently of *LSH1/LSH2*, whereas a large proportion of the developmental network depends on *LSH1/LSH2* activity for proper deployment during organogenesis. Considering the major function of *LSH1/LSH2*, it is perhaps surprising that the expression of these genes is predominantly associated with peripheral tissues of the nodule. We infer from this that *LSH1/LSH2* are likely to be inducing morphogens that move between cells to define development. Such morphogens are likely the hormonal programs that are entirely dependent on *LSH1/LSH2,* as well as perhaps mobile proteins such as SHR^21,46^.

Importantly, even severe root-like mutant organs do not acquire canonical lateral-root transcriptional programs, indicating that identity loss reflects regression toward a primary-root-associated ground state rather than trans-differentiation into lateral-root fate. More broadly, these findings suggest that organ identity regulators stabilize developmental states, and that identity loss may prolong precursor-like programs rather than directly converting tissues to alternative organ fates, as similarly observed in other developmental systems, including *Arabidopsis*^47,48^.

Our analyses highlight that cell identity is developmentally dynamic. Marker genes exhibit stage- and context-dependent behavior, and label transfer between datasets degrades when developmental stages are mismatched. These observations caution against treating marker genes as universally stable identifiers and suggest that many “cell types” inferred from single-cell datasets may instead represent spatially and developmentally restricted regulatory states shaped by developmental position, developmental time, and tissue context. With the rapid advancement of higher-resolution and higher-throughput spatial and single-cell technologies, future frameworks may benefit from interpreting cell identities as dynamic regulatory states rather than fixed categorical labels.

A nodule represents the integration of plant and bacterial developmental programs, and a key technological advance of this study is the demonstration that, through optimized probe design for plant and rhizobial transcripts, Xenium can simultaneously capture both host and bacterial gene expression while preserving spatial organization. Spatially resolved rhizobial gene expression across the interzone, nitrogen-fixation, and senescence regions suggests that bacterial functional states are aligned with host-defined developmental compartments. Consistent with this, co-expression analysis identified a shared host–microbe module containing the plant oxygen-buffering genes *MtLb1* and *MtLb2* together with bacterial oxygen-responsive respiration genes (*fixK1* and *fixNOP*), nitrogen fixation genes (*nifA* and *nifHDK*), and central carbon metabolism genes involved in dicarboxylate utilization and TCA-cycle function (*gltA*, *dctA*, *mdh*, *pckA*, and *dme*). This suggests that host oxygen buffering influences not only nitrogenase activity but also broader bacterial metabolic programs. Previous metabolic modelling demonstrated that oxygen limitation can constrain TCA cycle flux and 2-oxoglutarate production, thereby restricting ammonium assimilation and promoting nitrogen export to the host^49^. The coordinated expression of *gltA* with genes involved in dicarboxylate uptake and central carbon metabolism is consistent with this model and suggests that *gltA* enrichment may reflect an attempt to sustain TCA-cycle carbon flux within the low-oxygen nitrogen-fixation zone. The coordinated spatial expression of oxygen-buffering, respiration, nitrogen-fixation, and carbon-metabolism genes observed here highlights the ability of dual-species spatial transcriptomics to resolve integrated host–microbe physiological programs.

Our spatial transcriptome provides subcellular resolution not previously available, enabling functional interpretation of transcriptionally distinct cell states. One striking example is the separation of infected and uninfected populations within the nitrogen-fixation zone. While both plant and rhizobial gene expression support a primary role for nitrogen fixation in infected cells, uninfected cells exhibit distinct transcriptional programs consistent with specialized functions in nitrogen assimilation and nodule metabolism (Figure 2). The sharp spatial segregation of marker genes and signaling pathways between these populations further supports their functional differentiation. These findings suggest that uninfected nitrogen-fixation-zone cells are not simply cells lacking bacteria but represent a specialized cell type that contributes to the establishment and maintenance of an effective nitrogen-fixing organ.

In conclusion, our spatially resolved atlas reveals developmental insights that are inaccessible without spatial context. We demonstrate that nodule development emerges not from a single regulatory pathway but from the coordinated deployment of multiple concurrent developmental programs operating across distinct spatial domains. Spatial transcriptomics reveals how these programs are integrated during early organogenesis and subsequently partitioned into specialized cell-type-specific modules during maturation. Further, we show that nodule-specific cell types have no apparent counterparts in root development and that *LSH1/LSH2* are central regulators of these developmental programs, providing essential functions across diverse cell types during nodule formation. Consistent with the previously described role of *LSH* family genes in shoot development^23,50,51^, our findings highlight how common developmental regulators can be redeployed in distinct developmental contexts. The ability to resolve these concurrent regulatory states provides a framework for understanding how complex organs, such as the nodule, evolve and acquire unique identities, generating specialized cellular functions, including the capacity to accommodate nitrogen-fixing bacteria.

### Limitations of the study

Xenium profiles targeted gene panels rather than the whole transcriptome, constraining analysis to predefined genes and may limit detection of unanticipated regulatory components. Additionally, spatial resolution is dependent on segmentation strategy. While cell-wall–based and nucleus-anchored approaches yield consistent major cell states, each has biases in specific tissue contexts, particularly in densely packed meristematic regions or highly vacuolated cells. Future advances in whole-transcriptome spatial profiling, improved segmentation methods, and multimodal approaches integrating transcriptomics with chromatin accessibility, protein localization, and metabolic readouts will further refine the resolution of spatial regulatory networks. Genetic studies assume a function based on the specific genes mutated. Where genetic redundancy exists, as it does with all the genetic regulators studied here, we may be oversimplifying the functionality of these regulators. In a study such as this, one can always add additional time points or mutants, however, available resources limit our ability to saturate the study, and we have attempted to make best use of the resources that were available.

## RESOURCE AVAILABILITY

### Lead contact

Further information and requests for resources and data should be directed to and will be fulfilled by the Lead Contact, Min-Yao Jhu, and the corresponding author, Giles E. D. Oldroyd.

### Materials availability

This study did not generate unique materials or reagents. All materials used are commercially available or available through the *Medicago truncatula Tnt1* mutant database at Oklahoma State University.

### Data and code availability

- *Raw data and supporting information will be available from the date of publication by a peer-reviewed journal. During peer review, the data is available under controlled access. Reviewer access tokens are available from the corresponding author upon request*.
- Spatial transcriptomics data generated in this study have been deposited at the NCBI Gene Expression Omnibus (GEO) and will be publicly available as of the date of publication. These datasets include raw imaging data, decoded transcript coordinates, processed gene expression matrices, segmentation outputs, and associated metadata.
- All original code has been deposited at GitHub.

The Linux bash scripts used for relabeling and re-segmentation with Xenium Ranger are available at https://github.com/MinYaoJhu/Medicago_Xenium_relabeling_resegmentation.

The Python scripts for the cell wall and Cellpose-based cell segmentation pipeline are available at https://github.com/thiagomaf/PYcellsegmentation.

R scripts used for Xenium data analysis and visualisation are available at https://github.com/MinYaoJhu/Medicago_Rhizobia_Xenium_Analysis_Workflow.

R scripts for the re-analysis of published single-cell datasets are available at https://github.com/chongjing/scRNAseq_Medicago/.

An interactive Shiny application for exploring the processed Xenium data in RDS file format is available at https://github.com/MinYaoJhu/MedicagoXeniumSpatialViewer.

- Any additional information required to reanalyze the data reported in this paper is available from the lead contact upon request.

## ACKNOWLEDGMENTS

This work is supported by Gates Foundation and the UK Foreign, Commonwealth and Development Office (OPP1028264) and Gates Agricultural Innovations (INV-57461) known as the Enabling Nutrient Symbioses in Agriculture (ENSA) project. We gratefully acknowledge the 10x Genomics team—Ian Fiddes, Hank Tu, Yihui Zhu, Fallon Ratner, Nicola Cahill, and Rebecca Mitchell—for their assistance with custom Xenium panel design, data interpretation, and technical support in optimizing the workflow. We thank the Histopathology/ISH Core Facility at the Cancer Research UK Cambridge Institute—Elsa Santos, Julia Ponte, and Julia Jones—for technical support with H&E staining and bright-field slide scanning. We also thank the Cancer Research UK Cambridge Institute Genomics Core for support with multiple aspects of this study. We are particularly grateful to Victoria Winfield, Rachel Barnes, Alex Deamer, Ania Piskorz, and members of the CRUK single-cell team for performing Xenium workflows and generating raw data.

We thank Professor Uta Paszkowski and Professor Sebastian Schornack for valuable discussions and feedback on analytical strategy, and Eli Marable for constructive input that improved clarity and readability. We thank the CSC science support team, especially Phil Marshall and Kazuko Collins, for their help in maintaining growth conditions and preparing media for experiments, and the ENSA platform, led by Eleni Soumpourou, for providing DNA extraction services. Finally, we thank Pierre-Marc Delaux, Thomas Ott, Dugald Reid, Chun Liu, Anindya Kundu, Magdalini Tsitsikli, Bikash Raul, Beatrice Lace, Tatiana Vernie, Morgane Batzenschlager, Guofeng Zhang, Fabian van Beveren, Caspar Chater, and Carolina Isidra for sharing insights that support gene prioritization for probe design.

STPT image acquisition was supported by the IMAXT Cancer Grand Challenge grant (A24042 to Dario Bressan). Claire M. Mulvey and Dario Bressan are supported by the NIHR Cambridge Biomedical Research Centre (BRC-1215-20014). The views expressed are those of the authors and do not necessarily reflect those of the NIHR or the Department of Health and Social Care.

## AUTHOR CONTRIBUTIONS

M.-Y.J., G.E.O., and K.S. conceived the study. M.-Y.J. and C.L. performed sample harvesting and mutant genotyping. M.-Y.J. and J.H. optimized protocols and prepared FFPE samples and sections for Xenium. M.-Y.J., K.S., G.E.O., J.G., T.A.M., and C.X. contributed to gene selection and biological interpretation for Medicago probe design. P.P. provided biological expertise and interpretation on rhizobia gene selection for probe design. M.-Y.J. designed the Xenium Medicago and rhizobia probe panels with input and support from 10x Genomics. C.M.M. performed STPT imaging under the supervision of D.B., and D.B. performed 3D image alignment. M.-Y.J. performed Xenium preprocessing and spatial transcriptomics analyses. T.A.M. and M.-Y.J. developed and applied custom cell segmentation approaches. C.X. and M.-Y.J. performed single-cell reanalysis and label transfer. K.S. performed rhizobia infection visualization. M.-Y.J. wrote the manuscript with input from all authors. All authors reviewed, edited, and approved the final version.

## STAR★METHODS

### KEY RESOURCES TABLE

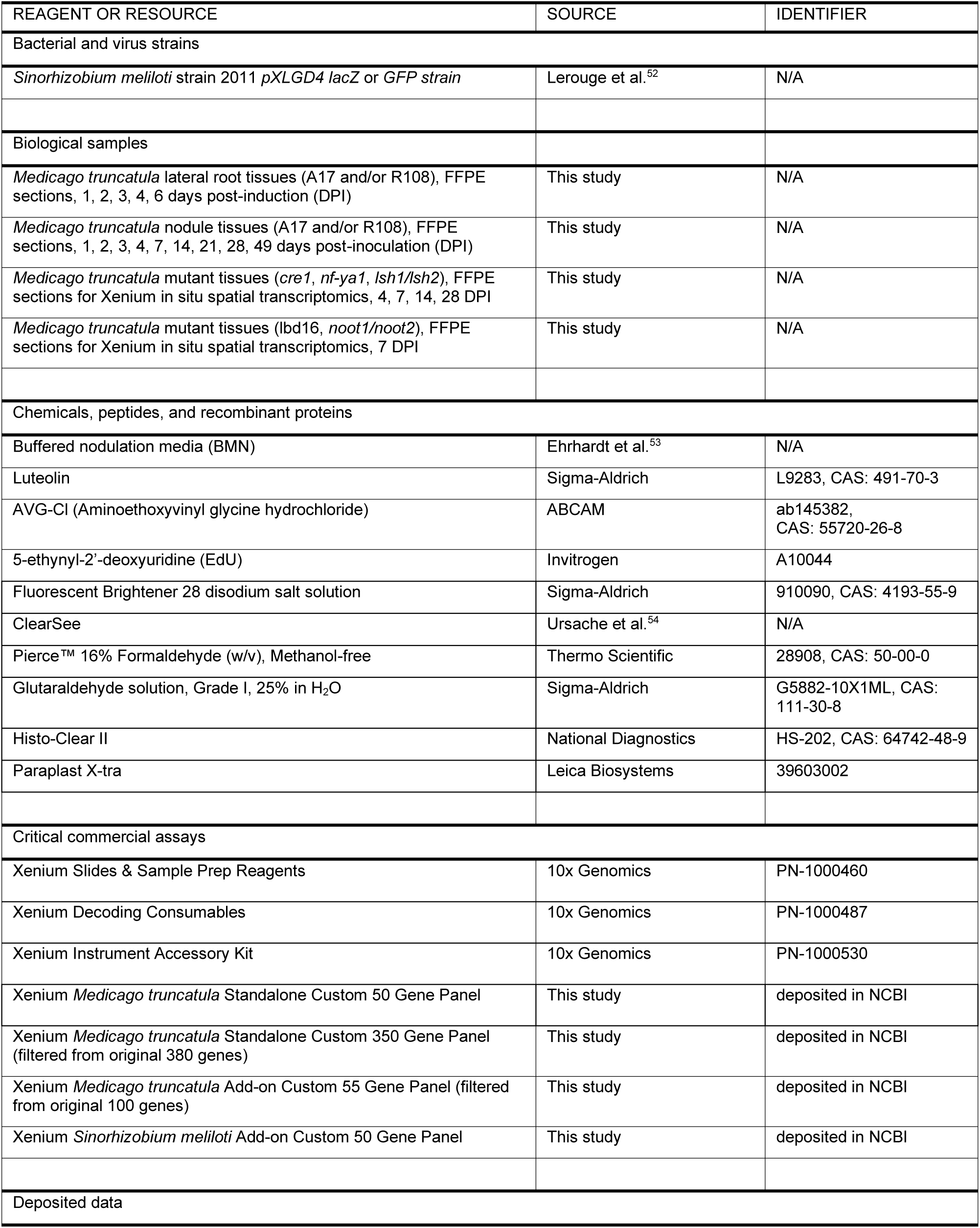
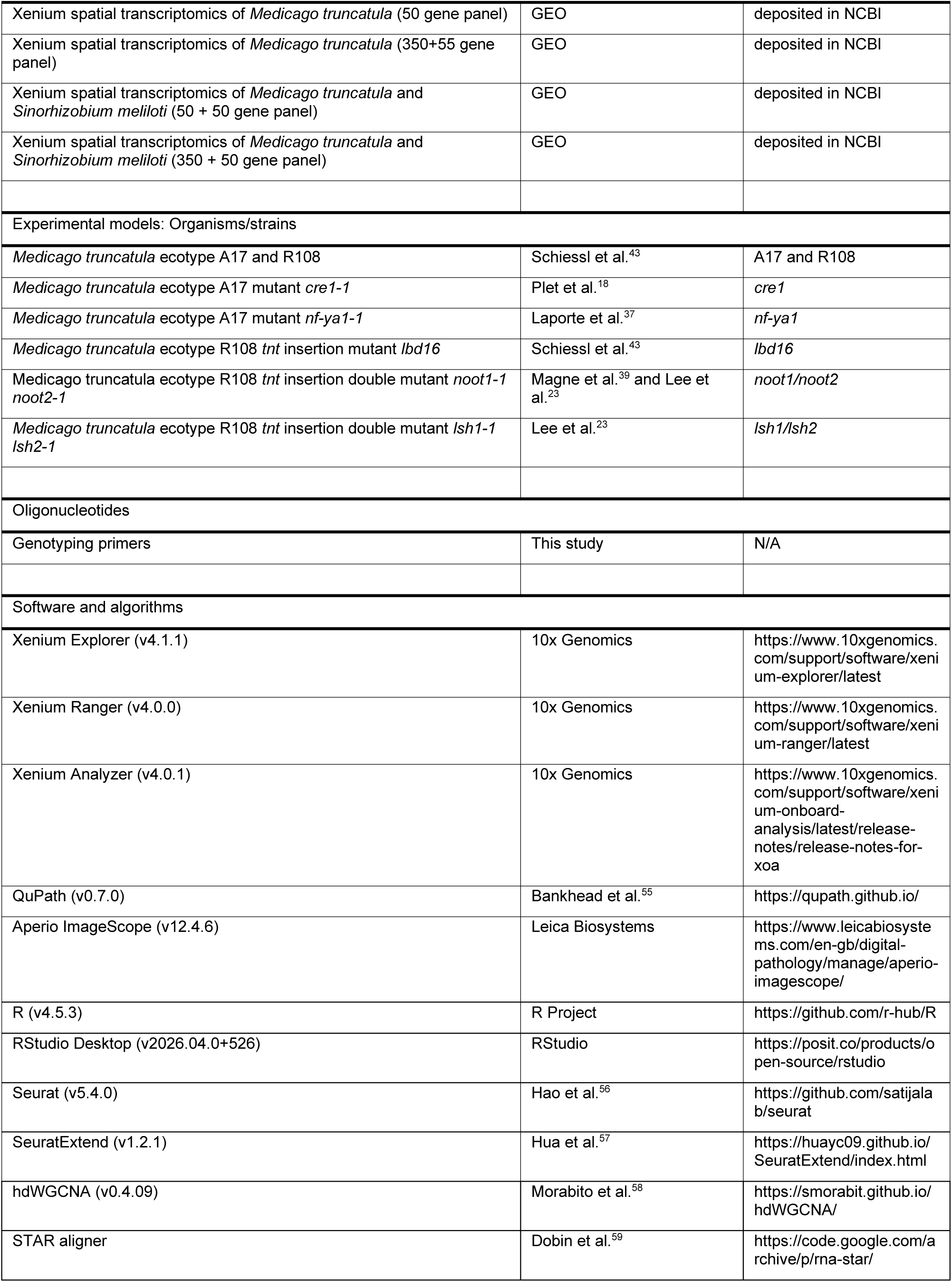
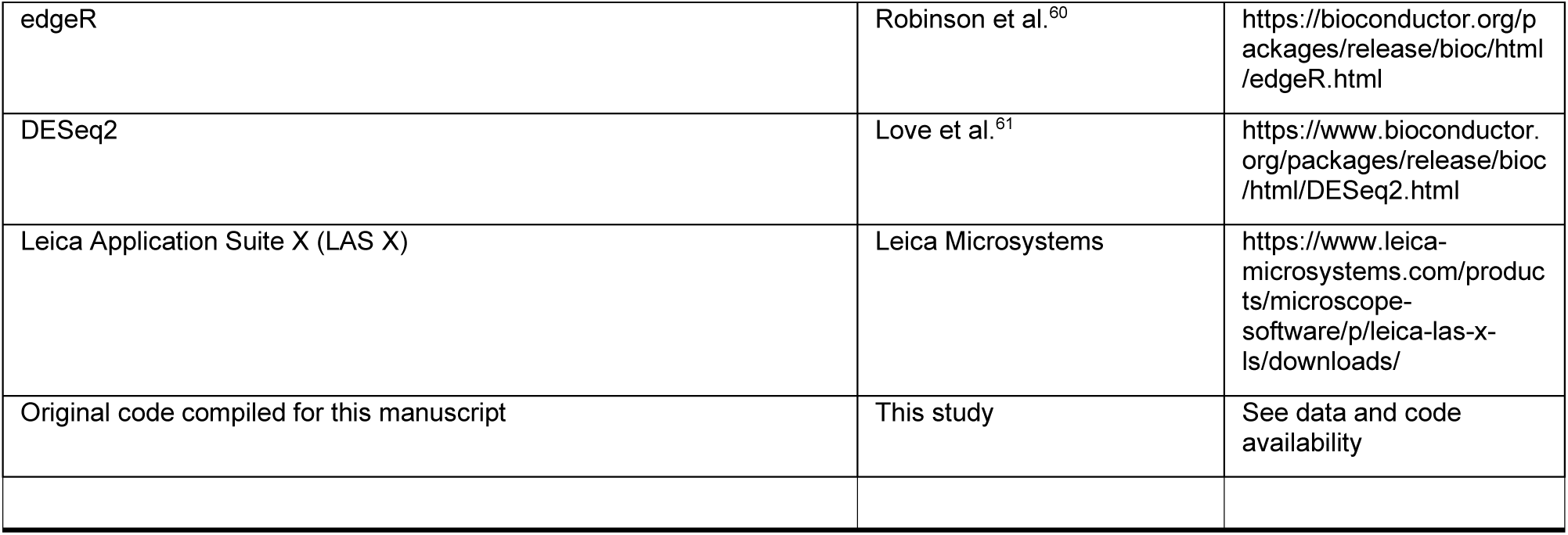

### EXPERIMENTAL MODEL AND STUDY PARTICIPANT DETAILS

#### Plant materials

*Medicago truncatula* ecotypes A17 and R108 were used as wild-type controls in this study. The mutant lines used have either the A17 or R108 genetic background, depending on their origin. The *nf-ya1* and *cre1* mutants (A17 background) were generated by ethyl methanesulfonate (EMS) mutagenesis and described previously ^37,38^. The *lsh1/lsh2* double mutant^23^, *noot1/noot2*^39^, and *lbd16* mutant^10^ (R108 background) are *Tnt1* retrotransposon insertion lines obtained from the *Tnt1* Retrotransposon Mutant Collection (Oklahoma State University, Stillwater, USA).

#### Bacterial strains

Two *Sinorhizobium meliloti* strains, Sm2011, were used in this study. The Sm2011–GFP strain, expressing a constitutive green fluorescent protein (GFP) marker, was employed for spot inoculation experiments to enable visualization of infection sites under a fluorescence dissecting microscope prior to tissue harvesting. This allowed verification of successful inoculation and accurate localization of early infection for sample collection. For spray inoculation experiments designed to capture mature nodules, the Sm2011–LacZ strain was used instead. The LacZ reporter minimizes background fluorescence and prevents interference with downstream Xenium *in situ* fluorescence detection. Both bacterial strains were cultured under identical conditions at 28 °C with shaking at 200 rpm, as described below, and diluted to the target optical density before inoculation.

### METHOD DETAILS

#### *Medicago* seed scarification and sterilization

*Medicago truncatula* seeds were scarified using concentrated sulfuric acid to remove the impermeable seed coat. Approximately 100 seeds were placed in 50 ml Falcon tubes, and concentrated H₂SO₄ was added under a fume hood until the seeds were fully submerged. Seeds were incubated for 4 min in acid, with gentle swirling every 30 s. When seed coats showed visible spotting, the acid was carefully removed, and seeds were rinsed six times with cold sterile distilled water to eliminate all acid residues.

Following scarification, seeds were placed on ice and transferred to a laminar flow hood for surface sterilization. Seeds were incubated in 0.75% (v/v) sodium hypochlorite for 10 min with gentle shaking, then rinsed six times with sterile distilled water. Sterilized seeds were kept in the dark in sterile water at room temperature on a shaker for 1 h to allow hydration. After soaking, the seeds were wrapped in aluminum foil and stored at 4 °C for 3 days for stratification.

For germination, stratified seeds were transferred onto sterile Petri dishes lined with moist filter paper, sealed with Parafilm, wrapped in foil, and incubated in the dark at 21 °C for at least 24 h. After 1–2 days, seeds typically showed radicle emergence and were ready for transfer to growth media or seed pouches for subsequent experiments.

#### Lateral root induction by gravity stimulation

Lateral root formation in *Medicago truncatula* was induced through gravity stimulation to achieve synchronized primordium development. Germinated seedlings were grown vertically on buffered nodulation medium (BNM) plates supplemented with phosphate (+P) but lacking nitrogen (−N) for 2 days under controlled growth conditions (21 °C, 16 h light/8 h dark photoperiod). After 2 days of vertical growth, plates were rotated 135° clockwise and incubated overnight to induce gravitropic bending of the primary root.

Control seedlings were marked at the root tip at the time of rotation but maintained in a vertical position throughout the experiment. Root samples were collected at 1, 2, 3, 4 and 5 days post-rotation (DPR) for fixation and spatial transcriptomic analysis. For each time point, only the bending region of the primary root was excised for downstream processing. The harvested root sections were inspected under a dissecting microscope to confirm the presence of lateral root primordia from 2 DPR onward, ensuring accurate capture of developing primordia for subsequent *in situ* profiling.

#### Spot inoculation for early nodule primordium induction

To capture early developmental stages of nodule primordia, *Medicago truncatula* seedlings were transferred to BNM medium without nitrogen (−N) supplemented with phosphate (+P) and 1 µM aminoethoxyvinylglycine (AVG) to inhibit ethylene biosynthesis. Seedlings were grown under controlled conditions (21 °C, 16 h light/8 h dark photoperiod) for 2 days before inoculation.

*Sinorhizobium meliloti* strain Sm2011 was used for inoculation. Bacteria were first cultured in tryptone–yeast extract (TY) medium and then subcultured 1 day prior to inoculation in minimal medium supplemented with 3 μM luteolin to activate symbiotic gene expression. Cultures were grown for ∼16 h at 28 °C with shaking until reaching an OD₆₀₀ of ∼1.0, and subsequently diluted to a final concentration of OD₆₀₀ = 0.025 in fresh Fahraeus (FP) medium.

For spot inoculation, 1 µl of the diluted bacterial suspension was carefully placed using a pipette onto the susceptible zone of each root, identified as the region where root hairs first emerge. The inoculation site was marked by punching the adjacent filter paper to facilitate accurate tracking. Root segments were harvested at 1, 2, 3, 4 and 7 days post-inoculation (DPI) for histological examination and spatial transcriptomic analysis.

#### Spray inoculation for mature nodule development

To obtain mature nodules, *Medicago truncatula* seedlings were grown under conditions optimised for sustained symbiotic development. Germinated seedlings were first transferred to BMM−N+P medium plates and grown vertically at 21 °C for 3 days. Seedlings were then transferred to seed pouches (CYG™ Grow Pouch, blue color, Mega International), which are light-opaque to minimize root illumination.

For transfer, a pre-made slit was gently opened in the pouch paper using sterilized forceps, and each seedling was carefully inserted so that the radicle extended into the moist pouch layer while the hypocotyl and cotyledons remained above the pouch surface. After all seedlings were transferred, two seed pouches were taped together face-to-face to completely block light exposure to the root system while allowing the shoot to remain illuminated. Seedlings were grown in the pouches for an additional 3 days at 21 °C, supplied with BMM−N + P liquid medium to maintain hydration.

*Sinorhizobium meliloti* strain Sm2011 was prepared for inoculation by initial culture in TY medium, followed by subculture 1 day prior to inoculation in TY medium supplemented with 3 mM luteolin to activate bacterial symbiotic gene expression. Cultures were grown overnight (∼16 h) until reaching an OD₆₀₀ of ∼1.0, and then diluted to a final concentration of OD₆₀₀ = 0.05 in fresh BMM−N + P medium.

For spray inoculation, the upper portion of each seed pouch was opened, and 2–3 ml of the bacterial suspension was evenly sprayed along the root surface using a syringe fitted with a nasal spray adaptor. Pouches were placed horizontally for 1 h to ensure even bacterial contact before being returned to a vertical position supported by bamboo seed pouch holders. Plants were maintained under standard growth conditions for the remainder of the experiment. Root nodules were harvested at 14, 21 and 28 days post-inoculation (DPI) for histological and spatial transcriptomic analyses.

#### Spatial transcriptomic sample preparation for Xenium *in situ*

Fresh *Medicago truncatula* nodules and root segments were harvested at defined developmental stages and processed as FFPE sections for Xenium *in situ* profiling. Step-by-step protocols are published^36^.

Briefly, tissues were fixed in freshly prepared 4% (w/v) paraformaldehyde and 0.25% (v/v) glutaraldehyde in 1× PBS (pH ∼7.4). Samples were submerged at a ≥1:10 tissue: fixative ratio, vacuum-infiltrated on ice (∼500 mm Hg, 20 min, four cycles), and incubated overnight at 4 °C (12–16 h). Dehydration at room temperature proceeded through an ethanol series (10%, 30%, 50%, 70%, 90%; 30 min each), followed by 100% ethanol for 1 h (×3). Samples were then transferred to fresh 100% ethanol and held overnight at 4 °C.

Clearing and infiltration used graded ethanol:Histo-Clear (3:1, 1:1, 1:3; 1 h each) followed by 100% Histo-Clear (×3, 1 h each). Paraplast X-tra (melted 58–60 °C) was introduced as 1:3 paraplast:Histo-Clear overnight (60 °C), then 1:2, 1:1, and 3:1 (3 h each), followed by 100% paraplast overnight with one additional overnight change in fresh paraplast. Tissues were embedded in pre-warmed molds, oriented to capture central (infection, interzone, fixation) zones of nodules, and stored at 4 °C.

Sections (5–10 µm) were cut on a rotary microtome. Initial sections were placed on standard slides to confirm orientation and the presence of the region of interest under a light microscope. Collection sections were floated briefly (5–10 s) on a 45 °C Milli-Q water bath and mounted onto Xenium slides (10x Genomics), ensuring placement within the capture area with sufficient spacing for multiple sections if required. Slides air-dried for 30 min at room temperature, baked 3 h at 42 °C, and desiccated overnight at room temperature. For deparaffinization, slides were baked 2 h at 60 °C, equilibrated to room temperature (∼10 min), then passed through xylene (10 min ×2), 100% ethanol (3 min ×2), 96% ethanol (3 min ×2), 70% ethanol (3 min), and nuclease-free water (20 s).

For Xenium chemistry, custom probe panels were reconstituted and prepared according to the manufacturer’s instructions (10x Genomics, Xenium *In situ* User Guide), with associated *.json* files uploaded to the instrument before the run. Padlock-type probes (two 20-nt RNA-binding arms) were heat-denatured (95 °C, 2 min), chilled on ice, combined with Probe Hybridization Buffer, applied at 500 µL per well, and incubated overnight on a pre-equilibrated thermocycler (Xenium V1 program). Decrosslinking, hybridization, ligation, rolling-circle amplification, autofluorescence quenching, and DAPI staining were performed according to the Xenium FFPE workflow, with minor adjustments for plant tissues.

Slides were loaded onto the Xenium Analyzer, and decoding reagents (Bottles 1–4 and Modules A/B) were prepared and run according to the manufacturer’s instructions. Regions of interest encompassing full nodule cross-sections and matched roots were selected using nuclear and autofluorescence channels. The instrument performed iterative hybridization–imaging–probe-removal cycles to generate optical barcodes and assign transcript identities. Per-ROI outputs (decoded transcripts, images, and QC report) were collected for downstream analysis.

Post-run, quencher removal (10 min in sodium hydrosulfite solution, followed by water rinses) allowed optional H&E staining and imaging for histological context. Detailed protocols for plant-compatible Xenium *in situ* workflow are published (Jhu et al., 2025).

#### Post-Xenium confocal imaging

Following Xenium *in situ* profiling, tissue sections were imaged using a Leica SP8 confocal microscope equipped with a white light laser and spectral detection. To improve single-cell segmentation in plant tissues, which often display irregular polygonal geometries, cell wall autofluorescence was leveraged to define cellular boundaries. Lambda-lambda spectral scans were first performed to characterize tissue-specific autofluorescence and determine optimal excitation and emission settings. Fluorescence lifetime imaging microscopy (FLIM) was then used to capture fluorescence decay profiles, allowing separation of overlapping signals based on lifetime characteristics. Derived lifetime parameters were subsequently applied for tau gating and tau separation to suppress background autofluorescence and enhance cell wall contrast in full-section images. The resulting images were used to generate cell boundary masks for spatial transcriptomic segmentation.

#### 3D Xenium sample preparation and H&E imaging data acquisition

The FFPE block was trimmed superficially to expose only the outer edge of the tissue. Serial sections were then taken at 10 μm intervals until the tissue was almost exhausted. The ribbon of serial sections (approximately 60 in number) was split into 4 roughly even ribbons, and each ribbon was placed on a pre-cleaned tray. The sections were counted, and the ribbons were kept in order so that the individual number of each section remained trackable. Three sections were removed from the full serial set, one at the start of the set (section 1), one in the middle (approx. section 30), and one at the end (approx. section 60). These were separated by placing the ribbon on a cutting mat and cutting out the individual sections with a razor blade. These were floated on a water bath at 45 °C for 5-10 seconds, collected onto a normal slide, and checked for orientation and quality under a microscope. Once deemed acceptable, the full serial set was collected, and the normal slide containing section 1 was used to collect from section 2 onwards. The first sections were collected onto normal slides until a section containing a fair representation of the nodule was reached, around section 10. From this point, sections were collected onto normal slides with individual sections cut out at roughly even intervals and collected onto Xenium slides. Each time a Xenium section was due to be collected, the immediately previous section was checked for quality under a microscope. The potential Xenium sections were also inspected by eye on the cutting mat; these checks allowed the best sections to be selected for Xenium, as quality varied slightly across the serial set. 8 Xenium sections were collected across 2 slides, with 4-6 sections in between. When the intermediate point was reached, the sections were collected onto the initial checking slide containing the mid-point section, so it was included in the serial set. The final approximately 10 sections were collected onto normal slides.

Normal slides were baked for 1 hour at 60 °C before being H&E stained on a multistainer, then scanned on a brightfield scanner. Xenium slides were air-dried for 30 minutes before being baked for 3 hours at 42 °C. They were then stored in a desiccator until the day of the Xenium run.

#### 3D phenotyping with Serial Two Photon Tomography

To prepare *Medicago* root nodules for serial two-photon tomography (STPT), the samples were embedded in 8 % (w/v) Type 1 agarose (Cat. No. A6013, Sigma) prepared in 50 mM phosphate buffer, pH 7.4. The agarose solution was melted and poured into a 2 cm³ mold (Peel-A-Way Embedding molds, Cat No. E6032, Sigma), on ice. The nodule was immersed in the molten agarose at the correct orientation, and the block was allowed to harden. 20 ml of 1X embedding polymer reagent was prepared according to the manufacturer’s instructions in a fume hood (Imbed 100S reagent, TissueVision, Cambridge, MA, USA), before being degassed for 30 minutes. Once solidified, the agarose block was placed in the embedding polymer solution for 72 hours at 4°C to allow the polymer to penetrate throughout the block. The block was then baked for 6 hours at 40°C to promote cross-linking of the polymer solution. The polymerized block was stored at 4°C in 50 mM phosphate buffer, pH 7.4, supplemented with 0.01 % sodium azide, until imaging.

Serial two-photon tomography imaging was performed using a TissueCyte® 1000 instrument, a high-speed multiphoton microscope with integrated vibratome for whole-mount serial sectioning and block-face imaging of a defined tissue volume (TissueVision, Cambridge, MA, USA). A series of 2D XY mosaic images was taken across the surface of the tissue (16x objective; resolution 500nm/pixel), followed by physical vibratome sectioning with a ceramic blade to remove the imaged tissue and to create a new surface for a subsequent round of imaging^62^. 150 sections of 15 µm thickness in the z-plane were sectioned at a speed of 0.1 mm/sec, 70 Hz frequency. A single imaging plane was acquired for each 15 µm physical section, with an excitation wavelength of 900nm (2-beam Chameleon 2-photon femtosecond laser, Coherent Discovery), equipped with filters centered at 450nm, 520nm, 630nm and 660nm. The tiled STPT images were stitched in 2D and aligned in 3D to create a reconstructed image of the nodule using the image analysis workflow described below and elsewhere^63^.

#### Visualization of rhizobial infection by β-galactosidase (x-gal) staining

For the detection of β-galactosidase activity, root tissues inoculated with Sm2011 rhizobia expressing the lacZ reporter gene (ProHemA:LACZ) were first rinsed in 50 mM phosphate buffer (pH 7.2) and subsequently fixed in 2.5% glutaraldehyde using vacuum infiltration for 15 min, followed by incubation at room temperature for 1 h. After fixation, samples were washed three times in Z-buffer (100 mM phosphate buffer pH 7.0, 10 mM KCl, 1 mM MgCl₂). Staining was performed in X-Gal solution prepared in Z-buffer supplemented with 5 mM potassium ferricyanide, 5 mM potassium ferrocyanide, and 5-bromo-4-chloro-3-indolyl β-D-galactopyranoside (blue X-Gal, Sigma-Aldrich, Darmstadt, Germany). Samples were incubated at 28°C for 6–12 h to allow color development, then washed three times with water. Following staining, nodules and roots were dehydrated through a graded ethanol series and stored in 70% ethanol at 4°C. For whole-mount imaging, tissues were mounted in 70% ethanol on glass slides and visualized using a Keyence VHX-5000 digital microscope (Keyence Ltd., Milton Keynes, UK) with magnifications ranging from 20× to 1000×.

### QUANTIFICATION AND STATISTICAL ANALYSIS

#### Xenium Medicago truncatula probe gene selection and panel design

Targeted gene panels for Xenium in situ spatial transcriptomics were designed to capture the regulatory architecture underlying nodule organogenesis, spatial zonation, and cell identity in *Medicago truncatula*. Gene selection was guided by integration of published single-cell and single-nucleus RNA-seq datasets, together with time-course bulk RNA-seq profiles of nodule and lateral root development (including Schiessl et al.; Roy et al.; Pereira et al.; Ye et al., 2022), enabling prioritization of genes associated with nodulation signaling, hormone response pathways, transcriptional regulation, meristem function, infection processes, and nitrogen fixation, as well as genes shared with or diverging from lateral root developmental programs. This strategy ensured representation of both core regulatory modules and organ-specific transcriptional signatures.

Candidate genes were subjected to a series of constraints to balance biological relevance with probe design feasibility and detection robustness. Genes not detected in reference single-cell datasets were excluded to ensure baseline transcript detectability. Probe design feasibility was enforced by requiring a minimum of three probe-binding sites per gene; genes with fewer than three conserved probes were deprioritized unless supported by ecotype-specific probe design. To enable cross-ecotype comparability, orthologs between the A17 and R108 genomes were considered, and genes lacking confident ortholog assignments were assigned lower priority. Expression-level filtering was applied to avoid both low-signal and saturation-prone targets: genes with extremely low expression across single-cell datasets were deprioritized unless supported by differential expression in developmental time-course datasets, whereas genes with very high constitutive expression but lacking dynamic regulation were also deprioritized to preserve sensitivity to spatial and temporal variation. Within these constraints, genes with established or hypothesized roles in nodulation and organogenesis were prioritized to maximize interpretability.

Two Xenium panels were implemented in this study, consisting of an initial 50-gene panel for targeted optimization and a larger custom panel for atlas-scale profiling. The final *Medicago* panel used for analysis comprised 405 genes, including 350 genes derived from an original 380-gene custom panel and 55 genes incorporated from an additional 100-gene add-on panel. Gene inclusion in the final analysis set followed the 10x Genomics relabeling workflow, retaining genes that passed quality control and annotation consistency criteria to ensure accurate mapping between probe design and transcript identity. Genes not retained in the final analysis set were excluded from downstream analysis but remain part of the broader panel design and will be reported separately in future work.

Probe design was optimized to maximize transcript detection across ecotypes by incorporating both conserved probes targeting shared sequences and ecotype-specific probes where required. For each gene, total probe coverage reflected the combined contribution of conserved and ecotype-specific probes, enabling robust detection while maintaining specificity.

Comprehensive annotation of gene identity, functional classification, inferred or reported cell-type specificity, ortholog mapping between A17 and R108 ecotypes, and probe design metrics is deposited in NCBI. This table also documents panel composition across the 50-gene and 405-gene configurations and integrates prior functional knowledge to support downstream interpretation of spatial expression patterns.

#### Xenium rhizobia probe selection and panel design

A custom rhizobia Xenium panel was designed to capture bacterial symbiont *Sinorhizobium meliloti* transcriptional states across nodule developmental zones while maintaining compatibility with the *Medicago* Xenium panels used in this study. Because bacterial transcripts are highly abundant within infected cells and bacteroids, probe design was optimized in collaboration with 10x Genomics by reducing the number of probes per target gene relative to the plant panel, thereby minimizing signal saturation while preserving robust transcript detection.

As single-cell reference datasets are not available for rhizobia during symbiosis, candidate genes were selected from previously published laser-capture microdissection and zone-specific transcriptomic studies of *Medicago* nodules^64^. Priority was given to genes exhibiting spatially restricted expression patterns associated with bacterial differentiation, nitrogen fixation, metabolism, and developmental transitions across nodule zones. This strategy enabled the selection of markers expected to distinguish transcriptional states across the infection zone, interzone, nitrogen-fixation zone, and senescence-associated regions.

The final rhizobia panel consisted of 50 genes and was designed to be modularly combined with either the *Medicago* 50-gene optimization panel (50 plant + 50 rhizobia targets; 100 genes total) or the *Medicago* 350-gene atlas panel (350 plant + 50 rhizobia targets; 400 genes total). This design enabled direct integration of host and bacterial spatial transcriptomic information while maintaining compatibility across experimental configurations. Annotation of rhizobia gene ID, gene name, and panel composition is deposited in NCBI.

#### Xenium data relabeling

After acquisition, the raw Xenium *in situ* datasets were processed using Xenium Ranger (10x Genomics) according to the manufacturer’s guidelines. The relabeling of decoded transcripts was performed following the official *Xenium Ranger* tutorial (available on the 10x Genomics support website). To ensure consistent transcript identifiers across the 50-gene and 405-gene panels, the corresponding panel JSON files were edited to synchronize target names before executing the “xeniumranger relabel” function. This step ensured uniform gene annotation across all datasets for downstream integration and analysis. The Linux bash scripts used for relabeling with Xenium Ranger are available at [GitHub link 0].

#### Xenium nucleus and cell segmentation

For nucleus segmentation, the default DAPI-based segmentation generated by Xenium Analyzer (10x Genomics) was used. For nucleus-containing cell segmentation, the nucleus segmentation boundary file generated in the Xenium Analyzer output was applied directly, with the --expansion-distance parameter set to 5, thereby defining each cell as the nucleus plus a 5-µm radial expansion or until the expanding boundary encountered a neighboring cell.

For cell wall-based cell segmentation, to achieve cell wall–based segmentation, fluorescence images capturing intrinsic cell wall autofluorescence, confocal-derived cell wall signals, or post-Xenium Hematoxylin & Eosin (H&E) images were used as input for Cellpose (version 4.0.7), a deep learning–based segmentation algorithm [Add Ref]. Images were processed using Cellpose with the ‘cpsam’ model, which is optimized for cytoplasmic and cell boundary detection. Segmentation parameters were optimized for plant tissue characteristics: cell diameter estimates (typically 60-90 pixels, adjusted based on tissue type and image resolution) and minimum cell size threshold (15 pixels). To determine the optimal mask for each sample, segmentation was performed, and the results were compared to selecting the best-performing output. The images were first converted to grayscale before segmentation, and for multi-dimensional image stacks, the maximum-intensity projection was applied along the z-axis.

The segmentation pipeline accepts images in the OME-TIFF format and extracts the spatial metadata, including the micrometers per pixel (MPP) value from the images to provide accurate scaling. Segmentation masks are created as labelled images, where each cell or nucleus is provided with a unique integer ID. These masks are used to assign transcripts to cells, where the cell ID of each cell containing a transcript is identified by determining the segmented region that contains each transcript’s spatial location. Detailed documentation of the scripts and a tutorial for cell segmentation and image processing can be found at https://github.com/thiagomaf/PYcellsegmentation. The resulting segmentation mask (.tiff) was then imported into Xenium Ranger using the import-segmentation function for re-segmentation.

Because post-Xenium images and Xenium transcript data have different spatial dimensions and coordinate systems, image alignment was performed to ensure accurate transcript reassignment. Alignment between the Xenium transcript data and the corresponding histological or confocal images was achieved using Xenium Explorer, following the official 10x Genomics tutorials. Briefly, confocal and H&E images were first converted to pyramidal, tiled OME-TIFF format for compatibility with Xenium Explorer. Image registration was then performed by manually placing at least 20 key points on corresponding anatomical landmarks between the Xenium image and the post-Xenium confocal or H&E image. Alignment accuracy was verified by inspecting the nuclei segmentation overlay. Once satisfactory alignment was achieved, the built-in registration function in Xenium Explorer was used to compute a transformation matrix aligning the post-Xenium image to the Xenium coordinate system. This transformation matrix was subsequently exported and used as input to Xenium Ranger during re-segmentation.

#### Post-Xenium image alignment and transformation

Following image import and registration, datasets containing both nucleus segmentation (from the default Xenium output) and cell segmentation (from Cellpose-derived masks) were processed using the xeniumranger import-segmentation function to ensure that all images, segmentation masks, and transcript coordinates were spatially co-registered within a unified coordinate system. Cell masks generated with diameter parameters of 60 and 90 were used for both confocal and H&E images, each aligned separately to the corresponding Xenium dataset. Alignment quality was evaluated by comparing the percentage of transcripts assigned to cells (transcript-in-cell ratio) and through manual inspection of segmentation boundaries in Xenium Explorer. The segmentation output demonstrated the highest transcript-in-cell ratio and most accurate visual alignment was selected for downstream analyses. The Linux bash scripts used for re-segmentation with Xenium Ranger are available at https://github.com/MinYaoJhu/Medicago_Xenium_relabeling_resegmentation.

#### Xenium in situ data preprocessing, integration and clustering

Xenium *in situ* transcriptomic data from 40 runs of the 405-gene panel were processed using R (v4.5.1) and Seurat (v5.3.1). For each run, relabeled or resegmentated outputs from the Xenium outs directory were imported using LoadXenium, which loads cell-level counts, molecule coordinates, cell centroids, and segmentation polygons. Only molecules/transcripts with a base-calling quality value ≥20 were retained (mols.qv.threshold = 20). To ensure global uniqueness after merging, cell barcodes and field-of-view (FOV) identifiers were prefixed with sample IDs. For cross-sample Xenium integration and analysis, the nucleus, along with a 5-µm radial expansion boundary used for spatial polygon plots, was explicitly set to the Xenium cell segmentation polygons.

Across all 40 datasets, we identified the intersection of detected genes and restricted analyses to this common feature set. After reconstructing the Xenium assay for each sample using only shared genes, all runs were merged into a single Seurat object while preserving associated images, molecule coordinates, and segmentation information. Quality control was performed using per-cell total UMI counts (nCount_Xenium) and detected genes (nFeature_Xenium). Cells with ≤1 UMI or ≤1 detected gene were removed. Negative-control probe counts were consistently near zero across all datasets, confirming minimal background signal and obviating the need for further filtering.

The QC-filtered object was normalized using log-normalization, and all expressed panel genes were used as variable features. Data was scaled and subjected to principal component analysis (PCA). Principal components explaining ≥1% of variance were retained for all downstream analyses. As an initial check, we computed uncorrected k-nearest neighbor graphs, clusters, and UMAP embeddings on the PCA space to assess sample-driven structure.

Batch correction across Xenium runs was performed via the IntegrateLayers framework in the Harmony integration at the embedding level, using sample identity as the batch variable. Neighbor graphs and clusters were recomputed on the Harmony embedding, and a batch-corrected UMAP was generated. Sample-level metadata, including genotype, tissue type, and developmental stage, were imported from an annotation file and joined to the combined Seurat object for subsequent stratified analyses.

We performed a systematic clustering-resolution sweep across Harmony embeddings (0.2–1.0). For each resolution, we computed silhouette scores on down-sampled Harmony-UMAP coordinates to identify the granularity that best balanced separation and stability. Marker-gene analysis was performed for selected resolutions using FindAllMarkers on the Xenium assay, after which we selected a final clustering resolution that maximized biological interpretability, silhouette performance, and consistency across replicates. The resulting cluster identities were stored as the final cluster assignment for all downstream analyses.

Final cluster marker genes were identified using Seurat’s FindAllMarkers function (Wilcoxon rank-sum test), restricting to positively enriched genes (only.pos = TRUE) detected in at least 5% of cells per cluster (min.pct = 0.05) without imposing a log-fold-change threshold. For visualization, the top three marker genes per cluster were selected by ranking genes within each cluster by decreasing average log2 fold change (avg_log2FC), using adjusted P value (p_val_adj) as a secondary tie-breaker. Duplicate genes that appeared as top markers across multiple clusters collapsed into unique entries for plotting.

To characterize cell-state structure across experimental factors, we quantified cluster composition across tissue type, genotype, and developmental stage. We conducted targeted comparisons for (i) A17 versus R108 (nodule and lateral root), (ii) wild type versus mutant genotypes within nodule tissue, and (iii) developmental time courses within A17 and R108. For each subset, key summary metrics (cluster proportions, gene-level signatures, and time-series transitions) were computed on the Harmony embedding and linked to known marker genes. The full analysis workflow, including all visualization code, is available in the accompanying GitHub repository: https://github.com/MinYaoJhu/Medicago_Xenium_analysis_workflow.

#### Pseudo-time analysis

Pseudo-time trajectories were inferred using Slingshot on the integrated Seurat object generated from A17 and R108 lateral-root and nodule spatial transcriptomic profiles. Lateral-root and nodule cells were analyzed separately according to the tissue_type annotation. Slingshot was run through SeuratExtend::RunSlingshot using PCA space when available, with Harmony space used as a fallback; Harmony UMAP coordinates were used only for visualization. Lineage-specific pseudo-time values were extracted from the Slingshot object, added to the Seurat metadata, and plotted on UMAP embeddings. Pseudo-time-dependent genes were identified with tradeSeq using Xenium count data. Genes detected in at least 20 cells with total counts of at least 20 were retained, ribosomal and mitochondrial genes were excluded, and negative-binomial generalized additive models were fitted with six knots using Slingshot pseudo-time and curve weights. Genes were ranked by association test, and the top 30 pseudo-time-associated genes were visualized as lineage-specific trend heatmaps. True time serves as an orthogonal reference, allowing pseudo-time trajectories to be interpreted as hypothesis-generating models that can be supported or refined by empirical temporal information.

#### Gene co-expression network analysis

Gene co-expression networks were constructed using hdWGCNA^58^, an R package designed for weighted gene co-expression network analysis in high-dimensional transcriptomic data, including single-cell and spatial modalities. Integrated spatial transcriptomic datasets (Xenium platform) were processed as described above. Cells were subset by genotype, tissue, and developmental stage, retaining only cells that passed quality-control filtering. For developmental network comparisons, nodule samples were grouped into early (4–7 DPI) and late (7–28 DPI) developmental stages. Lateral root samples (3–6 DPI) were analyzed as a comparative organ dataset. Given the sparsity inherent to spatially resolved transcriptomic data, we constructed meta-cells using k-nearest neighbor aggregation as implemented in hdWGCNA. Meta-cells were defined within genotype-specific subsets to preserve biological structure. Expression values were normalized prior to network inference.

Weighted gene co-expression networks were constructed using a soft-thresholding approach adapted from WGCNA^65^. To approximate scale-free topology, candidate soft-thresholding powers were evaluated using scale-free topology fit indices, and an optimal power was selected based on established criteria. An adjacency matrix was computed using the selected power and transformed into a Topological Overlap Matrix (TOM) to capture shared neighborhood connectivity^65^. Genes were hierarchically clustered based on TOM dissimilarity, and modules were identified using dynamic tree cutting. Each gene was assigned to a module represented by a unique color label. Module eigengenes (MEs) were calculated as the first principal component of scaled gene expression within each module.

To compare regulatory architecture across developmental stages and genotypes, the wild-type R108 nodule network was used as a reference. Intramodular connectivity (kME) was calculated as the correlation between individual gene expression profiles and their respective module eigengene ^65^. Genes were ranked by kME to identify highly connected hub genes within each module.

To identify genes most strongly associated with *LSH1*, Pearson correlations between *LSH1* expression and all genes in the wild-type R108 reference network were calculated using meta-cell expression profiles. Genes were ranked according to the absolute correlation coefficient with *LSH1*. The top 50 *LSH1*-associated genes were used to generate cross-condition correlation heatmaps, in which Pearson correlation coefficients between *LSH1* and each selected gene were visualized across developmental stages, genotypes, and organs. This approach enabled direct comparison of *LSH1*-associated regulatory relationships across conditions using an identical reference gene set.

To visualize developmental rewiring of *LSH1*-associated regulatory programs, signed gene co-expression networks were constructed using pairwise Pearson correlations among the top 45 *LSH1*-associated genes from the wild-type R108 reference network. Positive and negative correlations were represented as red and blue edges, respectively, and edge width reflected the absolute magnitude of the correlation coefficient. Node size was scaled according to the absolute correlation with *LSH1*, and node color indicated hdWGCNA module assignment. To facilitate direct comparison across developmental stages, genotypes, and organs, identical gene sets and node coordinates derived from the wild-type R108 reference network were maintained across all network visualizations.

Network architecture, module eigengene activity, and *LSH1*-associated correlation patterns were compared across developmental stages, genotypes, and organs to identify conserved and disrupted transcriptional programs. Changes in connectivity, module structure, and correlation patterns were interpreted as alterations in coordinated regulatory states associated with developmental transitions rather than direct regulatory interactions.

#### Re-analysis of published scRNA-seq datasets

Raw sequencing reads of single-cell and single-nucleus RNA-seq data from *Medicago truncatula* roots at the 0h, 24h, 48h, 96h and 14d after inoculation were downloaded from previously published studies ^26–28^. Sequencing reads of each sample were aligned to the reference genome MtrunA17r5.0 and processed using CellRanger (v7.2.0) ^66^. The raw expression count matrix was subjected to quality control and analyzed using Seurat (v5.1.0) ^67^. Specifically, cells with unique feature counts of 2,500 or more or 200 or fewer, and cells with>5% mitochondrial/chloroplast counts, were detected and filtered. Then, putative contamination from ambient RNA molecules was examined and corrected using ScCDC (v1.4)^68^. scDblFinder (v1.18.0)^69^ was applied to the corrected and normalized count matrix to detect and remove doublets. The two 14dpi samples were aggregated.

Since the 14d samples have distinct expression features, we only integrated 0h, 24h, 48h, 96h together, using two complementary approaches: one incorporating all expressed genes and another restricted to a conserved panel of 405 genes (that were used in spatial-transcriptome) for enhanced cross-sample comparability. After quality control, data were merged into a unified Seurat object, normalized and scaled, and batch-corrected via Harmony integration to mitigate sample-specific effects. Multi-resolution clustering was performed to identify cell populations, with cluster stability assessed using clustree^70^, and resolution 1.0 (all genes) or 0.4 (405 genes) selected for downstream analysis. Cell type annotation was assigned based on known marker genes ^26^, yielding distinct clusters corresponding to root tissue types, including epidermis, cortex, endodermis, stele, and nodule cells. More details on data processing are available at https://github.com/chongjing/scRNAseq_Medicago/.

### Declaration of generative AI and AI-assisted technologies in the manuscript preparation process

During the preparation of this work, the authors used Grammarly and ChatGPT 5.5 to assist with language editing, including improvements to grammar, clarity, and readability. The schematic diagram in Figure 7E and the schematic component of the bottom panel in the graphical abstract were created by the authors and subsequently refined with assistance from ChatGPT 5.5 for artistic improvement under careful author supervision and instruction. Following the use of these tools, the authors reviewed and edited all AI-assisted outputs as necessary and take full responsibility for the content of the published article.

