## Supplementary Figures for "4D spatial transcriptomics reveals nodule identity emerges through stacked parallel developmental programs"

[1]Crop Science Centre, Department of Plant Sciences, University of Cambridge, Cambridge, United Kingdom
[2]Cancer Research UK Cambridge Institute, University of Cambridge, Cambridge, United Kingdom
[3]Spatial Profiling and Annotation Centre of Excellence (SPACE), Cancer Research UK Cambridge Institute, University of Cambridge, Li Ka Shing Centre, Cambridge CB2 0RE, United Kingdom
[4]Department of Biology, University of Oxford, Oxford, United Kingdom
[5]Sainsbury Laboratory, University of Cambridge, United Kingdom
[6]The Donald Danforth Plant Science Center, Missouri, United States
[7]Present address: Novonesis, Gammel Venlighedsvej 14, 2970 Hørsholm, Denmark
[8]Present address: Plant Genetics, TUM School of Life Sciences, Technical University of Munich (TUM), Freising 85354, Germany
[9]Senior author
[10]Lead contacts

### SUPPLEMENTAL INFORMATION

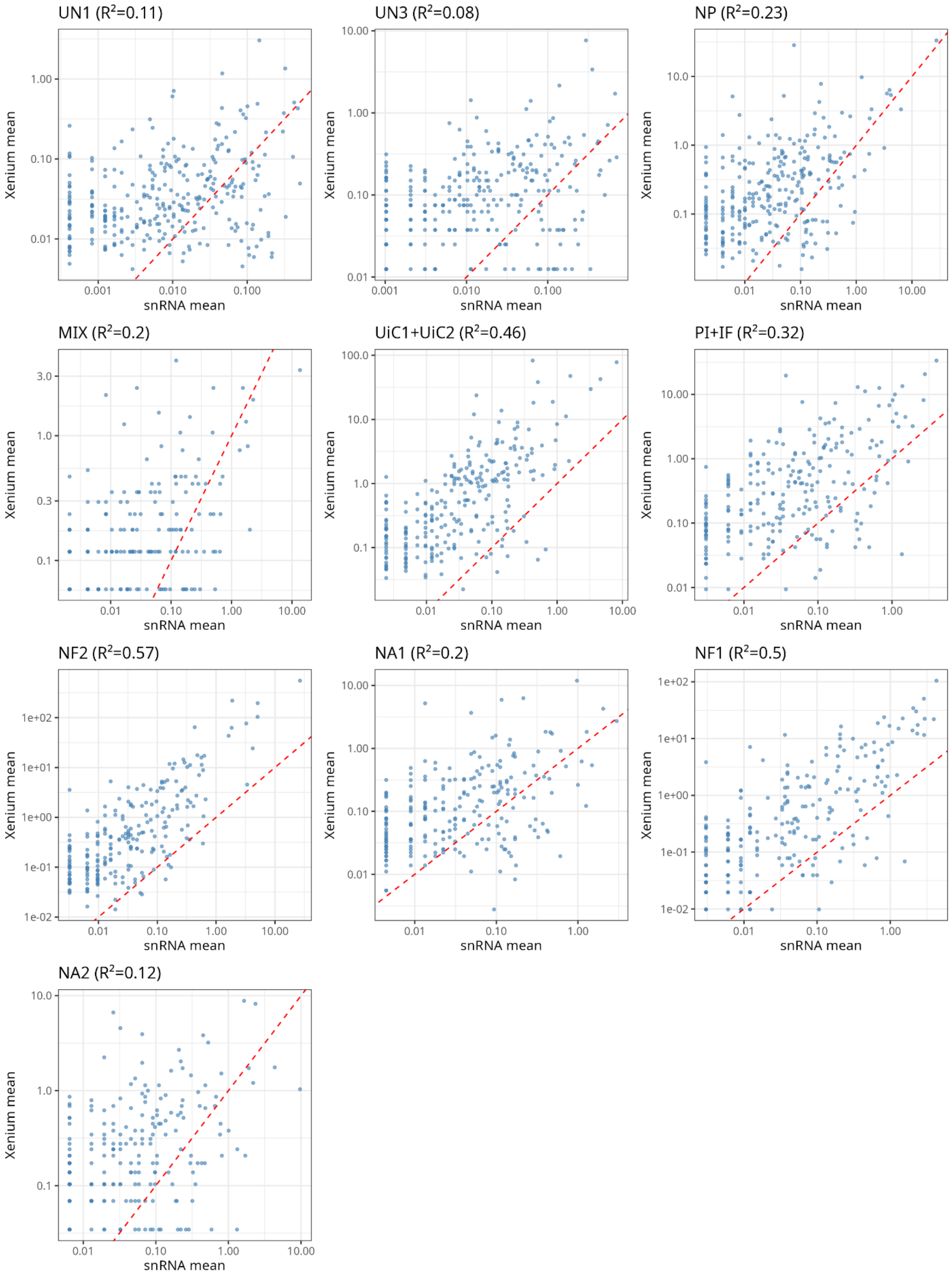

UN1 ($R^2$=0.11)
Xenium mean
1.00
0.10
0.01
0.001
0.010
0.100
snRNA mean
UN3 ($R^2$=0.08)
Xenium mean
10.00
1.00
0.10
0.01
0.001
0.010
0.100
snRNA mean
NP ($R^2$=0.23)
Xenium mean
10.0
1.0
0.1
0.01
0.10
1.00
10.00
snRNA mean
MIX ($R^2$=0.2)
Xenium mean
3.0
1.0
0.3
0.1
0.01
0.10
1.00
10.00
snRNA mean
UiC1+UiC2 ($R^2$=0.46)
Xenium mean
100.0
10.0
1.0
0.1
0.01
0.10
1.00
10.00
snRNA mean
PI+IF ($R^2$=0.32)
Xenium mean
10.00
1.00
0.10
0.01
0.01
0.10
1.00
snRNA mean
NF2 ($R^2$=0.57)
Xenium mean
1e+02
1e+01
1e+00
1e-01
1e-02
0.01
0.10
1.00
10.00
snRNA mean
NA1 ($R^2$=0.2)
Xenium mean
10.00
1.00
0.10
0.01
0.01
0.10
1.00
snRNA mean
NF1 ($R^2$=0.5)
Xenium mean
1e+02
1e+01
1e+00
1e-01
1e-02
0.01
0.10
1.00
snRNA mean
NA2 ($R^2$=0.12)
Xenium mean
10.0
1.0
0.1
0.01
0.10
1.00
10.00
snRNA mean

**Figure S1. Per–cell-type correlation between snRNA-seq and Xenium expression profiles following label transfer (cell-wall segmentation)**

Per–cell-type correlation between the 14-DPI snRNA-seq reference dataset and the integrated 405-gene Xenium spatial transcriptomic dataset following label transfer using cell-wall segmentation. For each transferred cell type, cells assigned the same label in the reference (snRNA-seq) and spatial (Xenium) datasets were aggregated, and mean gene expression was calculated across shared genes. Only genes detected in both datasets and expressed in both populations were included.

Scatter plots show log10-transformed mean expression values for snRNA-seq (x-axis) versus Xenium (y-axis). The dashed line indicates equality (y = x). Each panel corresponds to a single transferred cell type, and $R^2$ values were calculated using Pearson correlation. Cell types with fewer than 15 cells per dataset or fewer than 15 shared expressed genes were excluded.

Abbreviations: NA1/NA2 (nodule apex), PI (pre-infection), IF (infection), NF1/NF2 (nitrogen fixation), NP (nodule parenchyma), VA (vascular), UiC1/UiC2 (uninfected cells), UN1–3 (unknown), MIX (mixed identity).

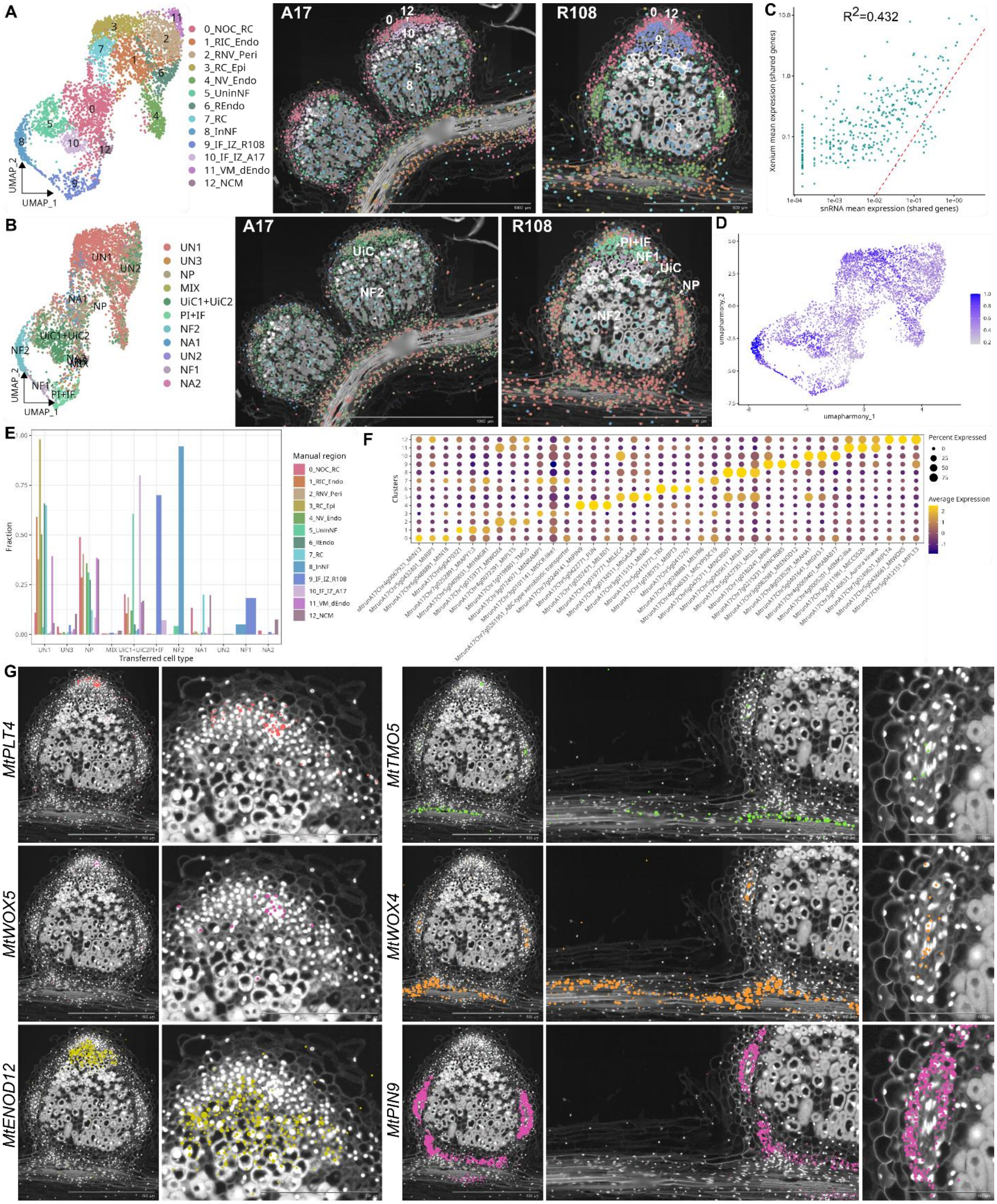


**Figure S2. Integrating spatial transcriptomics with single-cell data using nucleus-anchored segmentation in *Medicago* nodules**

(A) Left, UMAP of the nucleus-anchored spatial transcriptomic atlas of wild-type *Medicago truncatula* A17 and R108 nodules at 14 DPI, showing clustering based on spatial gene expression and histological context. Thirteen major cell types were identified: nodule outer cortex and root cortex (NOC_RC, cluster 0); root inner cortex/endodermis (RIC_Endo, cluster 1); root and nodule vasculature (pericycle; RNV_Peri, cluster

2); root cortex/epidermis (RC_Epi, cluster 3); nodule vasculature (endodermis; NV_Endo, cluster 4); uninfected cells in the nitrogen-fixation zone (UninNF, cluster 5); root endodermis (REndo, cluster 6); root cortex (RC, cluster 7); infected cells in the nitrogen-fixation zone (InNF, cluster 8); infection zone + interzone (R108) (IF_IZ_R108, cluster 9); infection zone + interzone (A17) (IF_IZ_A17, cluster 10); vasculature meristem/dividing endodermis (VM_dEndo, cluster 11); and nodule central meristem (NCM, cluster 12). Right, corresponding spatial maps showing localization of annotated cell types in representative A17 and R108 nodules.

(B) UMAP colored by labels transferred from a 14-DPI single-cell RNA-seq dataset, with corresponding spatial projections. Abbreviations: NA1/NA2 (nodule apex), PI (pre-infection), IF (infection), NF1/NF2 (nitrogen fixation), NP (nodule parenchyma), VA (vascular), UiC1/UiC2 (uninfected cells), UN1–3 (unknown), MIX (mixed identity).

(C) Correlation of normalized gene expression between single-cell RNA-seq and Xenium spatial transcriptomic datasets across shared genes. Each point represents one gene; the diagonal indicates equal expression.

(D) UMAP colored by label-transfer confidence scores, with darker blue indicating higher prediction confidence.

(E) Comparison between spatially defined clusters and transferred single-cell labels, illustrating correspondence between native spatial identities and single-cell annotations.

(F) Dot plot showing expression of marker genes defining the 13 spatial cell types identified in (A).

(G) Xenium *in situ* visualization of representative marker transcripts, including *Medicago truncatula* PLETHORA 4 (*MtPLT4*), WUSCHEL-RELATED HOMEOBOX 5 (*MtWOX5*), EARLY NODULIN 12 (*MtENOD12*), TARGET OF MONOPTEROS 5 (*MtTMO5*), WUSCHEL-RELATED HOMEOBOX 4 (*MtWOX4*), and PIN-FORMED 9 (*MtPIN9*), shown from whole-organ to cellular resolution in 14-DPI nodules. Nucleus-anchored segmentation improved detection of densely packed meristematic, vascular, and primary-root-associated populations, including low-abundance markers such as *WOX5*, whereas cell-wall segmentation more effectively resolved enlarged cortical and nitrogen-fixation–zone cells.

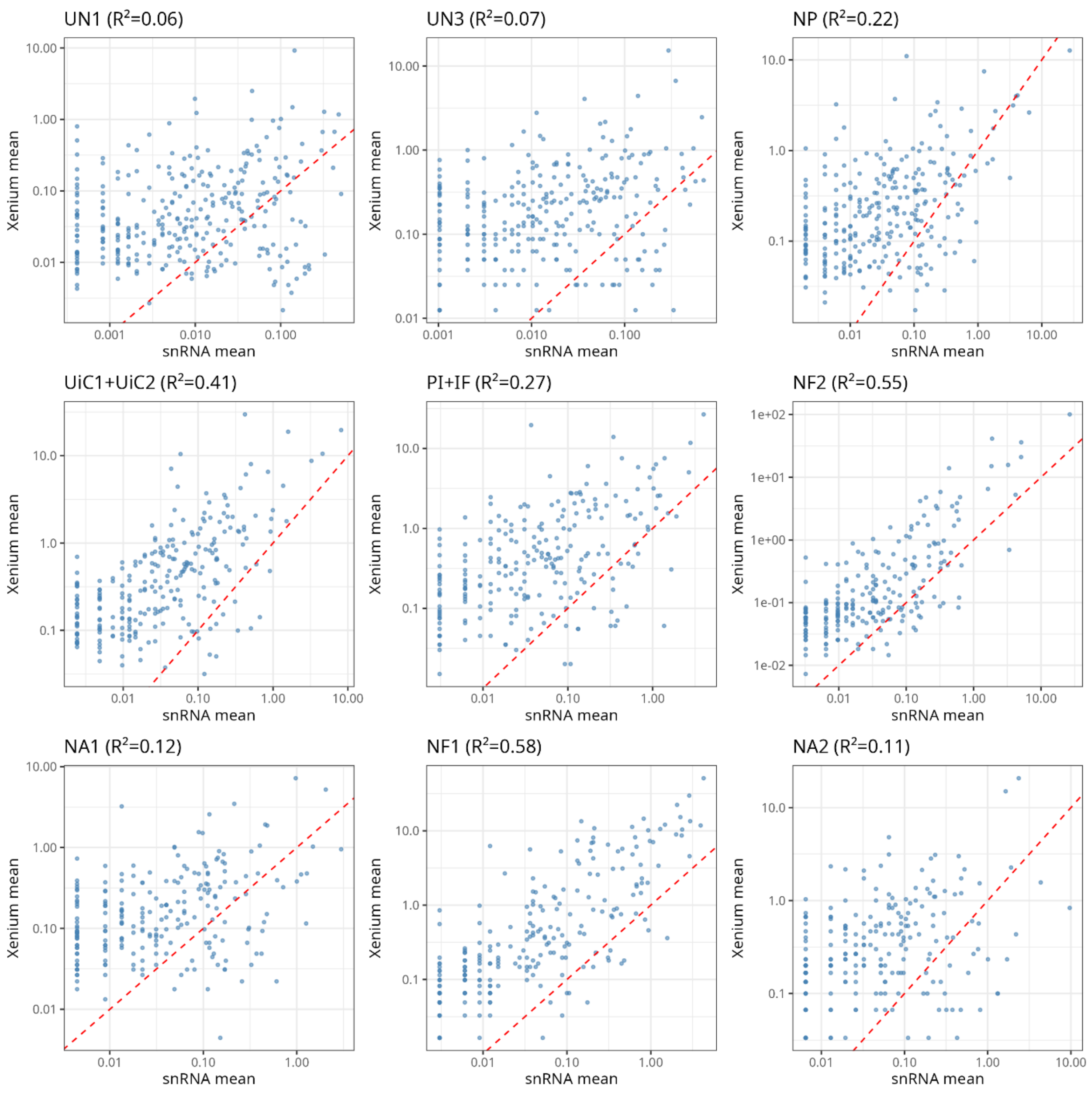


**Figure S3. Per–cell-type correlation between snRNA-seq and Xenium expression profiles following label transfer (nucleus-anchored segmentation)**

Per–cell-type correlation between the 14-DPI snRNA-seq reference dataset and the integrated 405-gene Xenium spatial transcriptomic dataset following label transfer using nucleus-anchored segmentation. Analysis was performed as in Figure S1.

For each transferred cell type, cells assigned the same label in the reference and spatial datasets were aggregated, and mean gene expression was calculated across shared genes. Only genes detected in both datasets and expressed in both populations were included.

Scatter plots show log10-transformed mean expression values for snRNA-seq (x-axis) versus Xenium (y-axis). The dashed line indicates equality (y = x). $R^2$ values were calculated using Pearson correlation. Cell types with fewer than 15 cells per dataset or fewer than 15 shared expressed genes were excluded.

Abbreviations are as defined in Figure S1. NA1/NA2 (nodule apex), PI (pre-infection), IF (infection), NF1/NF2 (nitrogen fixation), NP (nodule parenchyma), VA (vascular), UiC1/UiC2 (uninfected cells), UN1–3 (unknown), MIX (mixed identity).

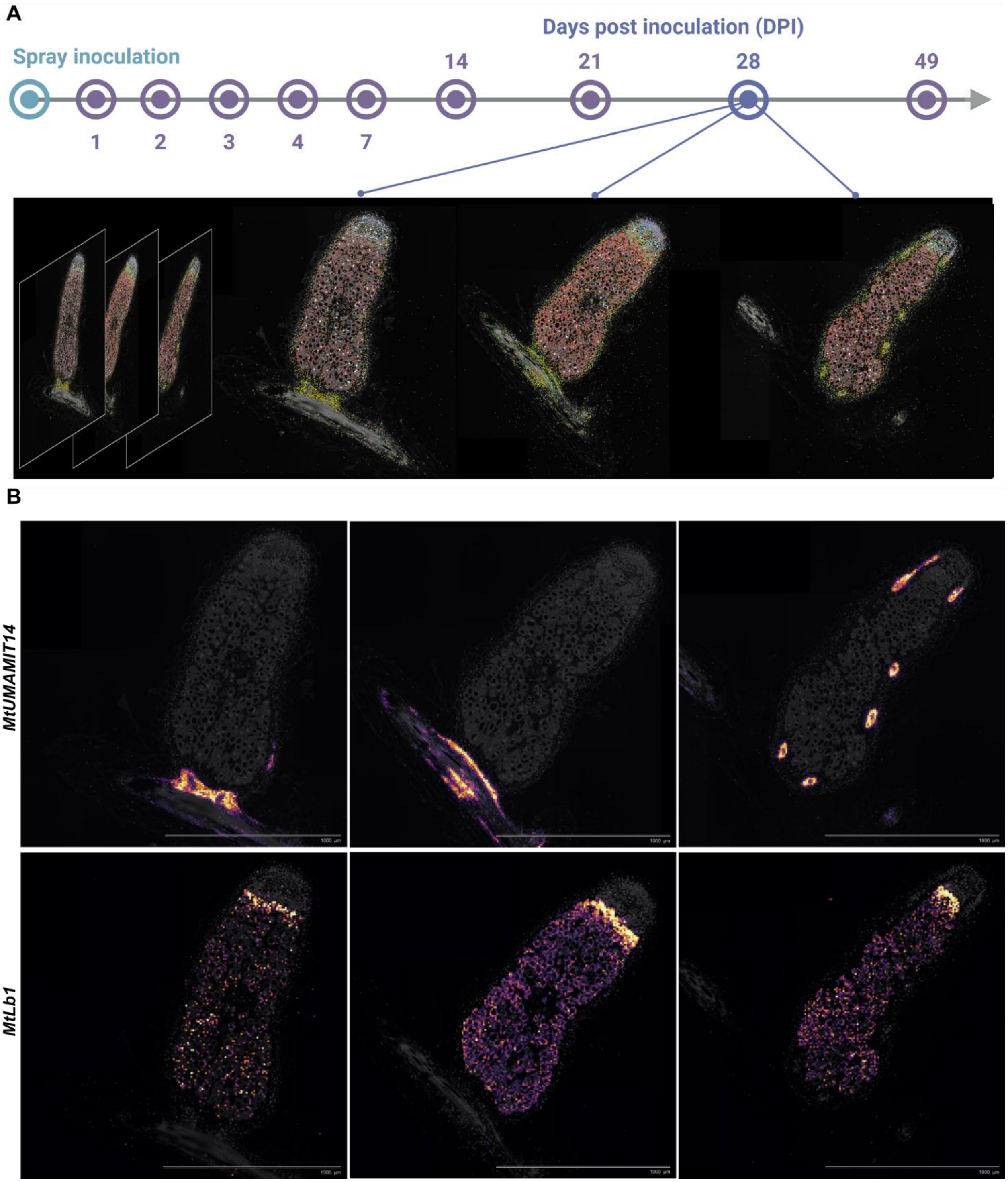


**Figure S4. Serial-section spatial transcriptomics reveals three-dimensional organisation of vascular and spatial gene gradients in mature nodules**

(A) Serial sections of a mature *Medicago truncatula* nodule (28 DPI) prepared at 10 µm, 8 µm, and 5 µm thickness. These matched sections enable reconstruction of late-stage nodule architecture and provide the basis for three-dimensional spatial alignment.

(B) Serial Xenium sections from a single 28-DPI nodule illustrate how two-dimensional views can obscure gene-expression patterns arising from three-dimensional tissue organisation. Top, *UMAMIT14*, a marker of nodule vasculature, shows strong expression in vascular strands that shift position across consecutive sections, reflecting the curvature of the vascular network. Bottom, *Leghemoglobin 1* (*Lb1*) exhibits a spatial gradient, with highest expression at the interzone–nitrogen-fixation boundary and progressively reduced signal through the fixation zone, while remaining confined to infected cells.

Together, these data demonstrate that apparent two-dimensional expression patterns often reflect underlying three-dimensional structure, highlighting the importance of multi-section or three-dimensional reconstruction for accurate interpretation of spatial gene expression in nodules.

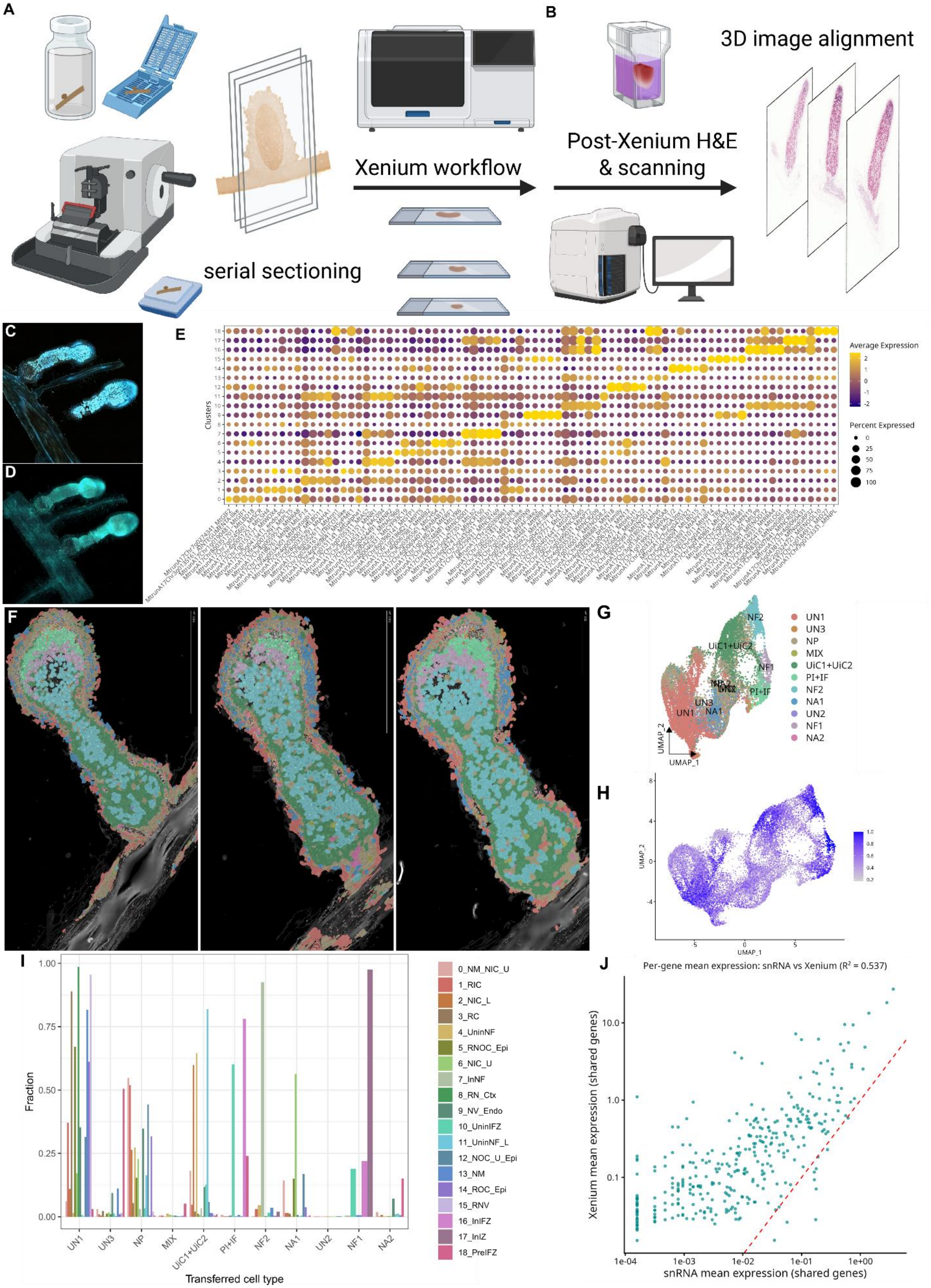
A
B
Xenium workflow
serial sectioning
Post-Xenium H&E
& scanning
3D image alignment
C
D
E
Clusters
Average Expression
Percent Expressed
0
25
50
75
100
F
G
UN1
UN3
NP
MIX
UiC1+UiC2
PI+IF
NF2
NA1
UN2
NF1
NA2
UMAP_1
UMAP_2
H
I
Fraction
Transferred cell type
0_NM_NIC_U
1_RIC
2_NIC_L
3_RC
4_UninNF
5_RNOC_Epi
6_NIC_U
7_InNF
8_RN_Ctx
9_NV_Endo
10_UninIFZ
11_UninNF_L
12_NOC_U_Epi
13_NM
14_ROC_Epi
15_RNV
16_InIFZ
17_InIZ
18_PreIFZ
J
Per-gene mean expression: snRNA vs Xenium (R² = 0.537)
Xenium mean expression (shared genes)
snRNA mean expression (shared genes)

**Figure S5. Three-dimensional spatial organization and cross-stage validation of 49-DPI spatial cell identities by marker expression and label transfer**

(A–B) Workflow for 3D spatial transcriptomic reconstruction of 49-DPI *Medicago truncatula* nodules using serial Xenium profiling, histological alignment. Nodules from the same stage were analyzed by Serial Two-Photon Tomography (STPT) imaging.

(C–D) Representative STPT sections and 3D renderings showing vascular organization and rhizobia-associated autofluorescence within mature nodules.

(E) Dot plot showing expression of representative top marker genes for major cell clusters identified in the 49-DPI 405-gene spatial transcriptomic dataset (Figure 2).

(F) Spatial projection of cell-type labels transferred from a 14-DPI single-cell RNA-seq dataset onto the 49-DPI spatial transcriptomic dataset. Abbreviations are as defined in Figure S2.

(G) UMAP of the 49-DPI spatial dataset colored by transferred 14-DPI single-cell labels. Abbreviations: NA1/NA2 (nodule apex), PI (pre-infection), IF (infection), NF1/NF2 (nitrogen fixation), NP (nodule parenchyma), VA (vascular), UiC1/UiC2 (uninfected cells), UN1–3 (unknown), MIX (mixed identity).

(H) UMAP showing label-transfer confidence scores, with darker blue indicating higher prediction confidence. Abbreviations are as defined in Figure 2 and S1.

(I) Comparison between spatially defined clusters and transferred single-cell labels, illustrating correspondence between native spatial identities and single-cell annotations.

(J) Correlation of normalized gene expression between single-cell RNA-seq and Xenium spatial transcriptomic datasets across shared genes. Each point represents one gene; the diagonal indicates equal expression.

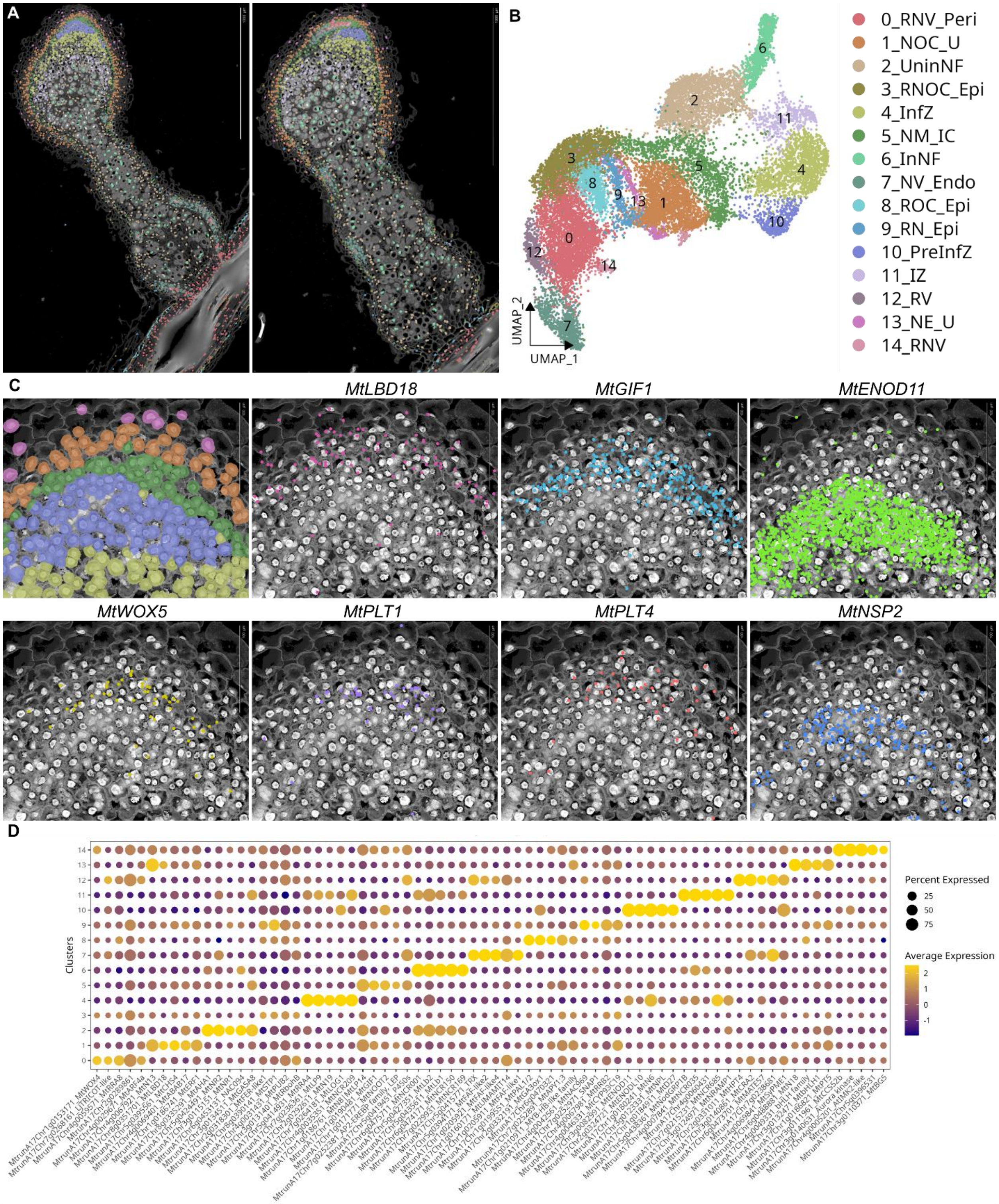


**Figure S6. Nucleus-anchored segmentation improves spatial resolution of apical nodule domains**

(A) Spatial projection of nucleus-anchored clustering for the 49-DPI spatial dataset, showing spatial distribution of cluster identities.

(B) UMAP embedding of nucleus-anchored clusters. Fifteen clusters were identified: root and nodule vasculature (pericycle; RNV_Peri, cluster 0); nodule outer cortex (upper; NOC_U, cluster 1); uninfected cells in the nitrogen-fixation zone (UninNF, cluster 2); root and nodule outer cortex + epidermis (RNOC_Epi, cluster 3); infection zone (InfZ, cluster 4); nodule meristem and inner cortex (NM_IC, cluster 5); infected cells in the nitrogen-fixation zone (InNF, cluster 6); nodule vasculature (endodermis; NV_Endo, cluster 7); root outer cortex + epidermis (ROC_Epi, cluster 8); root and nodule epidermis (RN_Epi, cluster 9); pre-infection zone (PreInfZ, cluster 10); interzone (IZ, cluster 11); root vasculature (RV, cluster 12); nodule epidermis (upper; NE_U, cluster 13); and root and nodule vasculature (RNV, cluster 14).

(C) High-resolution view of the apical meristem region. Nucleus-anchored segmentation resolves densely packed meristematic tissues into five spatially organized layers from apex to base: nodule epidermis (NE_U, cluster 13), upper outer cortex (NOC_U, cluster 1), nodule meristem and inner cortex (NM_IC, cluster 5), pre-infection zone (PreInfZ, cluster 10), and infection zone (InfZ, cluster 4). Spatial expression of representative marker genes—including *LOB DOMAIN-CONTAINING PROTEIN 18* (*MtLBD18*), *GRF-INTERACTING FACTOR 1* (*MtGIF1*), *EARLY NODULIN 11* (*MtENOD11*), *WUSCHEL-RELATED HOMEOBOX 5* (*MtWOX5*), *PLETHORA 1* (*MtPLT1*), *PLETHORA 4* (*MtPLT4*), and *NODULATION SIGNALING PATHWAY 2* (*MtNSP2*)—supports this stratified organization. Cell-wall and nucleus-based segmentation produced broadly concordant major cell identities, although nucleus-anchored segmentation improved detection of densely packed apical and vascular tissues, whereas cell-wall segmentation better resolved enlarged nitrogen-fixation and cortical cells.

(D) Dot plot showing expression of representative top three marker genes for major clusters identified in the 49-DPI dataset using nucleus-anchored segmentation.

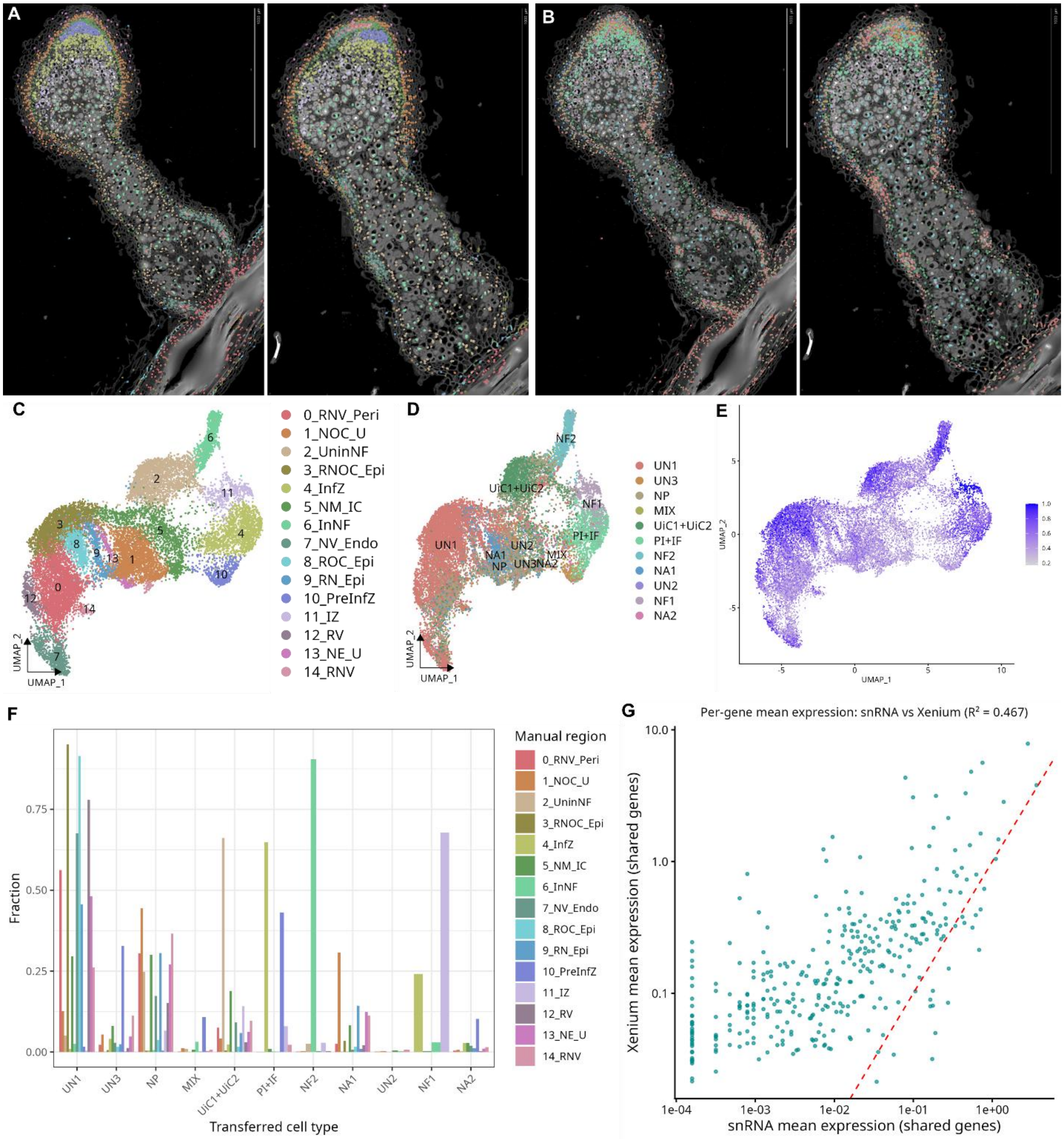


**Figure S7. Cross-stage validation of nucleus-anchored spatial cell identities at 49 DPI**

(A) Spatial projection of cluster identities defined from the 49-DPI Xenium dataset using nucleus-anchored segmentation.

(B) Spatial projection of cell-type labels transferred from a 14-DPI single-cell RNA-seq reference dataset onto the 49-DPI nucleus-anchored spatial dataset.

(C) UMAP embedding coloured by nucleus-anchored cluster identities. Cluster annotations correspond to those defined in Figure S6.

(D) UMAP embedding coloured by transferred single-cell RNA-seq labels. Major cell identities—including nodule apex (NA1, NA2), pre-infection (PI), infection zone (IF), nitrogen fixation (NF1, NF2), nodule parenchyma (NP), vascular tissue (VA), and uninfected cell populations (UiC1, UiC2)—remain clearly

separable following label transfer. UN1–UN3 denote clusters without a distinct marker signature; MIX indicates a cluster lacking strong identity enrichment.
(E) UMAP showing label-transfer confidence scores, with darker blue indicating higher prediction confidence. Confidence distributions are comparable to those observed for cell-wall segmentation (Figure S5).
(F) Comparison between native spatial clusters and transferred single-cell labels, illustrating concordance between nucleus-anchored spatial identities and single-cell annotations.
(G) Correlation of normalised gene expression between single-cell RNA-seq and nucleus-anchored Xenium spatial transcriptomic datasets across shared genes. Each point represents one gene; the diagonal indicates equal relative expression between platforms.

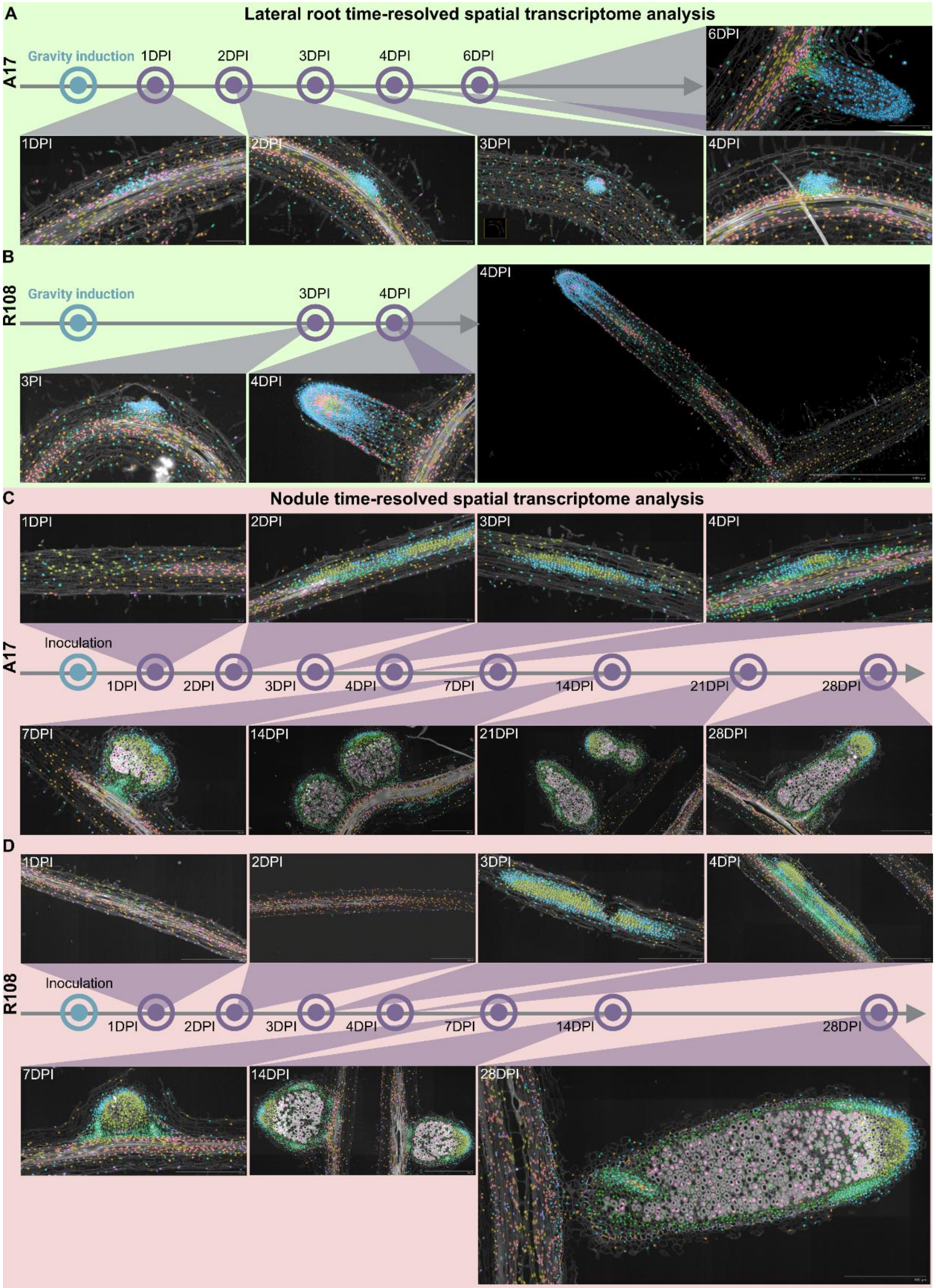


**Figure S8. Spatiotemporal organization of lateral root and nodule development across ecotypes**

(A–B) Time-resolved spatial transcriptomic maps of lateral roots following gravity induction, colored by shared UMAP-defined cluster identities.

(A) A17 lateral roots at 1, 2, 3, 4, and 6 days post-rotation (DPR).

(B) R108 lateral roots at 3 and 4 DPR.

These panels illustrate progressive spatial organization of lateral root tissues and conservation of major cell-type identities across ecotypes.

(C–D) Time-resolved spatial transcriptomic maps of nodules following rhizobial inoculation, colored by the same cluster identities as in (A–B), enabling direct comparison between nodule and lateral root developmental programs.

(C) A17 nodules at 1, 2, 3, 4, 7, 14, 21, and 28 days post inoculation (DPI).

(D) R108 nodules at 1, 2, 3, 4, 7, 14, and 28 DPI.

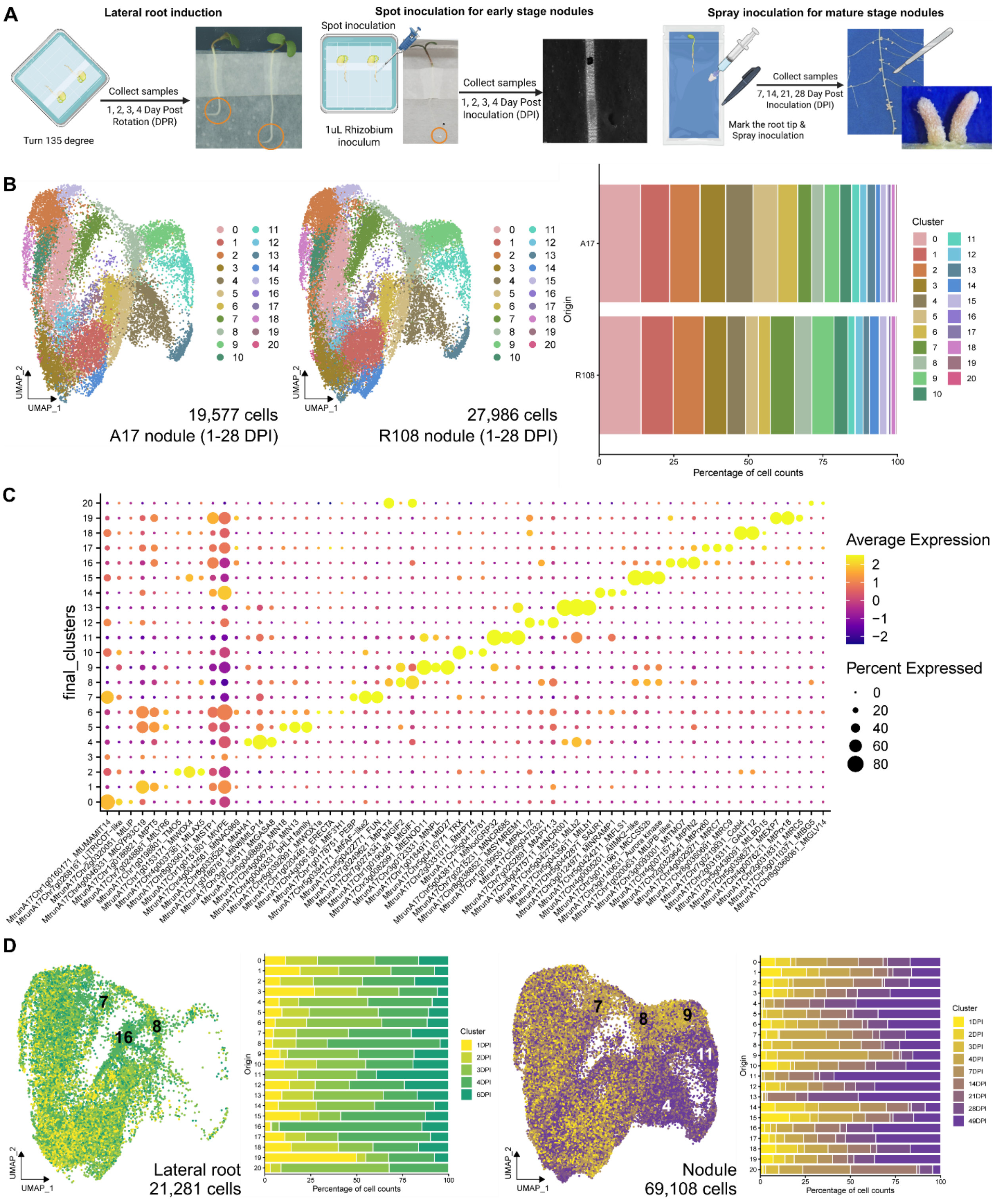


**Figure S9. Ecotype comparison of cell composition, marker definition, and organ temporal progression**

(A) Workflows for spatial transcriptomic sampling of lateral roots and nodules across developmental time.
(B) Ecotype-integrated UMAP comparison of Harmony-integrated Xenium data from wild-type A17 and R108, colored by the 21 cluster identities defined in Figure 3: cluster 0, Root endodermis (REndo); cluster 1, Root cortex (RCx); cluster 2, Root + nodule vasculature (stele) (RNV_St); cluster 3, Root epidermis +

cortex (REpiCx); cluster 4, Uninfected cells in the nitrogen-fixation zone (UninfNF); cluster 5, Root cortex + nodule upper outer cortex (RCx_UOC); cluster 6, Root cortex + nodule lower outer cortex (RCx_LOC); cluster 7, Root + nodule vasculature (endodermis) (RNV_Endo); cluster 8, Nodule + root meristem/primordium (NRMerPr); cluster 9, Pre-infection zone (PIF); cluster 10, Root + nodule pericycle (RNP_Pc); cluster 11, Infection zone (IF); cluster 12, Root inner cortex (RIC); cluster 13, Infected cells in the nitrogen-fixation zone (InfNF); cluster 14, Root epidermis + outer cortex (REpiOC); cluster 15, Root endodermis + pericycle (REndoPc); cluster 16, Lateral root tip cortex (LRTipCx); cluster 17, Lateral root cap (LRCap); cluster 18, Root vasculature (stele) (RV_St); cluster 19, Lateral root epidermis + cortex (elongation zone) (LR_EpiCx_EZ); cluster 20, Root meristem (RMer). Left, A17 cells; middle, R108 cells; right, bar plot showing proportional representation of each cluster in A17 and R108. Major spatial cell states are conserved across ecotypes, with quantitative differences in cluster composition.

(C) Marker-gene definition of spatial cell types. Dot plot showing expression of representative marker genes defining the 20 clusters identified in Figure 3 using nucleus-anchored segmentation. Dot size indicates the fraction of expressing cells, and color intensity represents average expression.

(D–E) Temporal progression of cell populations across developmental stages. UMAPs are colored by true developmental time with corresponding bar plots.

(D) Lateral-root cells (1–6 DPI), showing continuous temporal progression from early (yellow) to late stages (dark green).

(E) Early nodule development (1–7 DPI), colored from early (yellow) to later stages (purple), highlighting temporal dynamics during primordium formation and early differentiation.

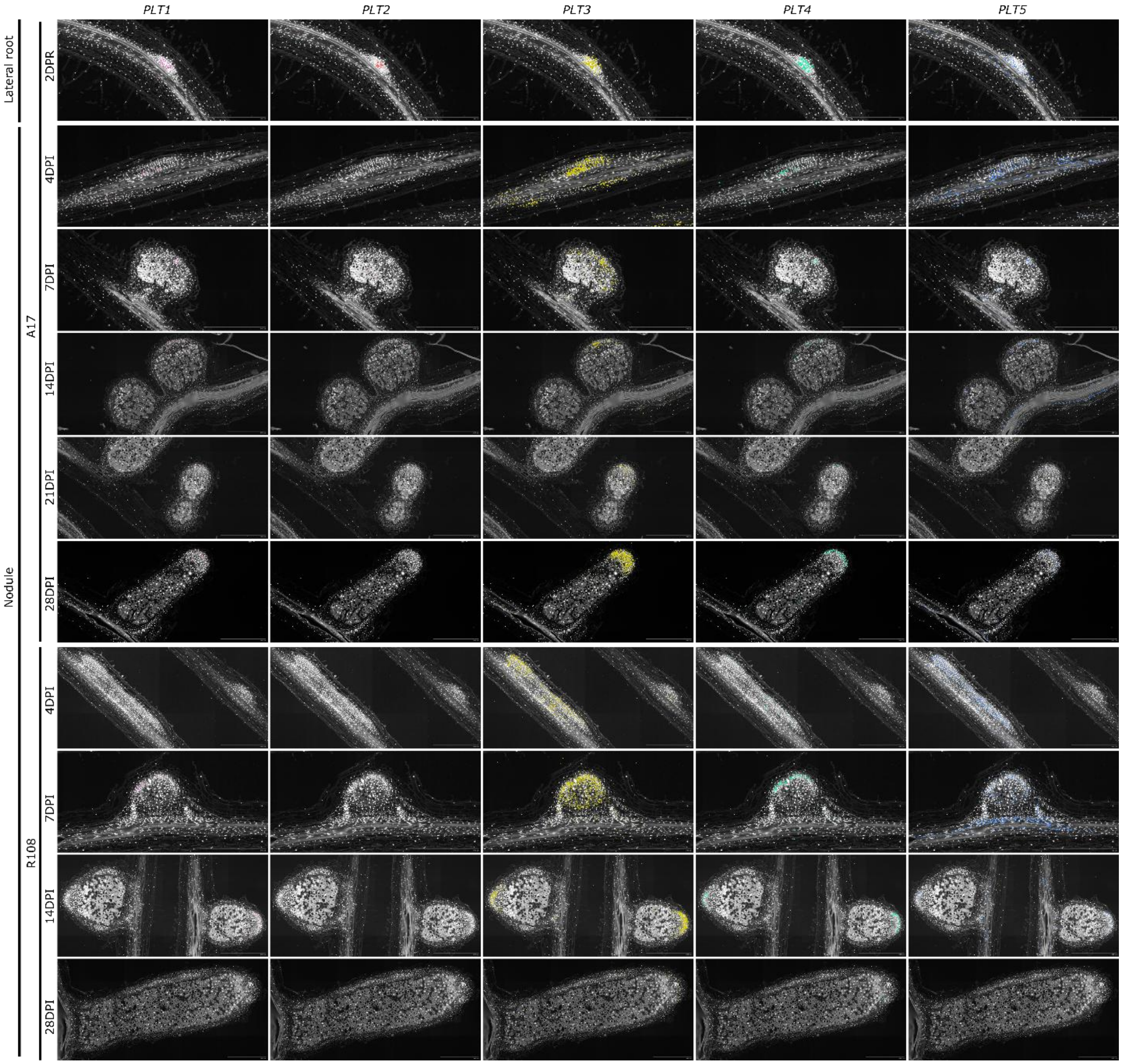


**Figure S10. Spatiotemporal expression dynamics of *PLETHORA* transcription factors during lateral root and nodule development**

Spatial transcriptomic maps showing the distribution of *PLETHORA* transcription factors *PLT1*, *PLT2*, *PLT3*, *PLT4*, and *PLT5* during lateral root and nodule development in *Medicago truncatula*. Transcript localization is visualized as colored spots overlaid on tissue sections. The first row shows a lateral root primordium at 2 days post-rotation (DPR), providing a reference for *PLT* expression during early lateral root development. Subsequent rows show nodule development in the A17 ecotype at 4, 7, 14, 21, and 28 days post inoculation (DPI), followed by corresponding stages in the R108 ecotype at 4, 7, 14, and 28 DPI. Across the developmental time course, *PLT1*–*PLT5* exhibit dynamic, stage-specific, and spatially restricted expression patterns.

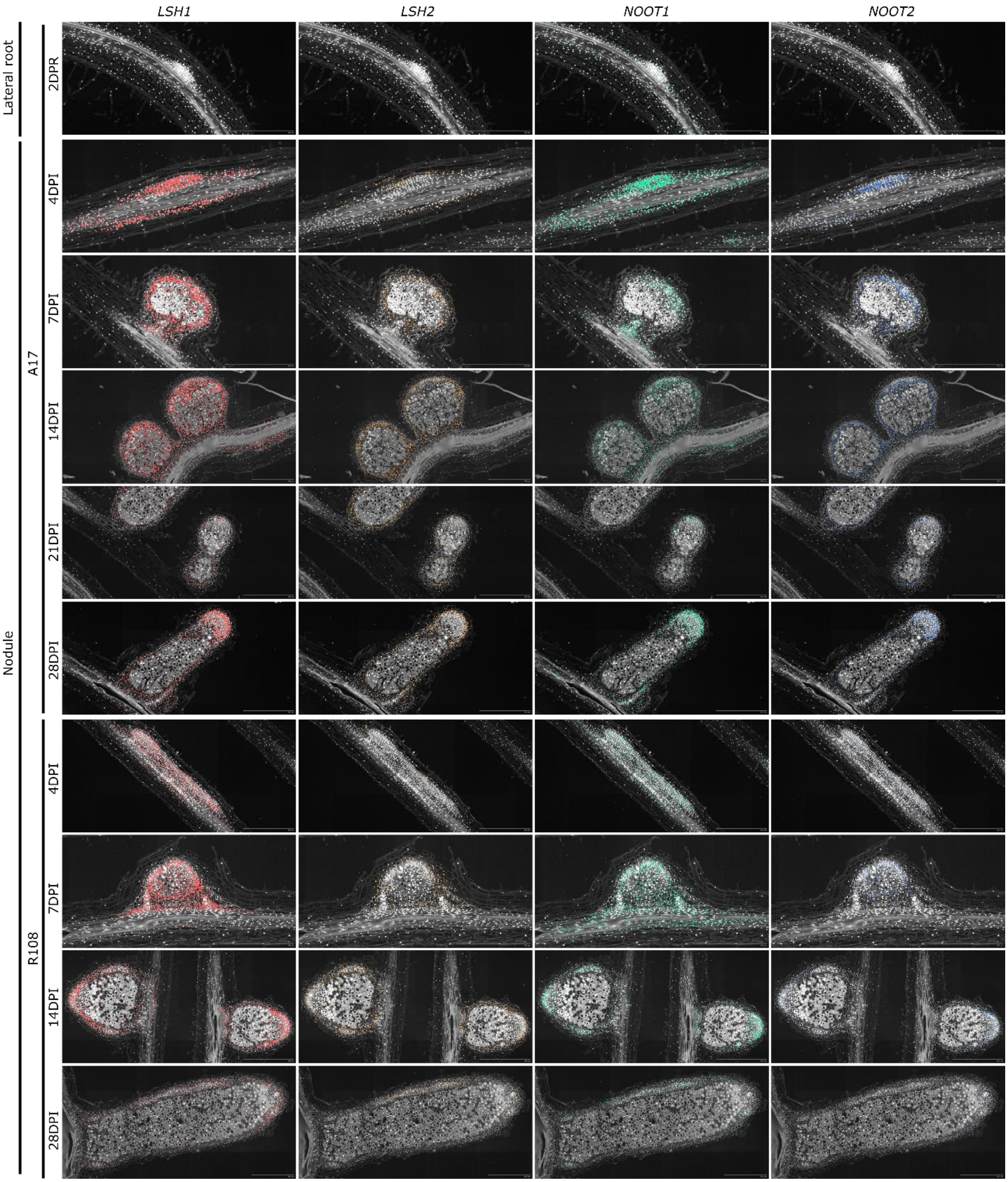


**Figure S11. Spatiotemporal expression dynamics of *LSH* and *NOOT* transcription factors during lateral root and nodule development**

Spatial transcriptomic maps show the distribution of key transcription factors involved in nodule identity and peripheral tissue regulation, including *LSH1*, *LSH2*, *NOOT1*, and *NOOT2*, during lateral root and nodule development in *Medicago truncatula*. Transcript localization is visualized as colored spots overlaid on tissue sections. The first row shows a lateral root primordium at 2 days post-rotation (DPR), providing a baseline reference for gene expression during early lateral root development. Subsequent rows show nodule development in the A17 ecotype at 4, 7, 14, 21, and 28 days post inoculation (DPI), followed by corresponding stages in the R108 ecotype at 4, 7, 14, and 28 DPI.

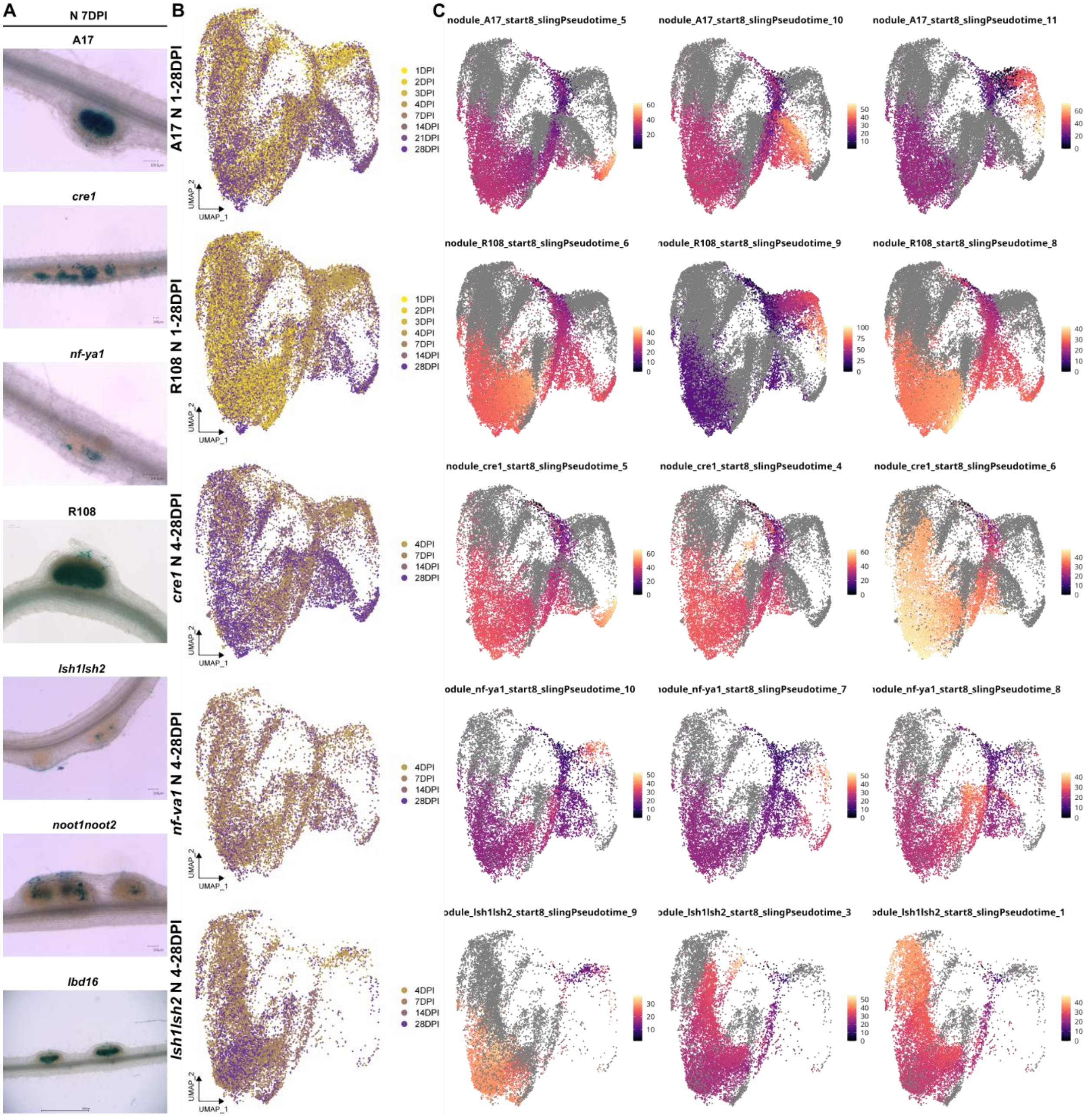


**Figure S12. Genotype-specific rhizobial infection patterns and developmental pseudo-time trajectories in nodulation mutants**

(A) Whole-mount X-Gal staining of *Medicago truncatula* roots and nodules inoculated with *Sinorhizobium meliloti* Sm2011 expressing a constitutive *lacZ* reporter (*ProHemA:lacZ*). Blue precipitate indicates β-galactosidase activity and marks rhizobial colonization and infection sites. All samples were imaged at 7 days post inoculation (DPI). Genotypes are arranged as follows (top to bottom): A17 wild type, *cre1*, *nf-ya1*, R108 wild type, *lsh1/lsh2*, *noot1/noot2*, and *lbd16*.

(B) True-time-colored UMAP embeddings for nodule datasets analyzed separately by genotype. From top to bottom: wild-type A17 nodules (1–28 DPI), wild-type R108 nodules (1–28 DPI), *cre1* nodules (4–28 DPI), *nf-ya1* nodules (4–28 DPI), and *lsh1/lsh2* nodules (4–28 DPI). Cells are colored by developmental stage, from early (yellow) to late (purple), illustrating temporal progression within each genetic background.

(C) Slingshot pseudo-time trajectories inferred independently for each genotype using the shared primordium/meristem population (NRMerPr; cluster 8) as the trajectory root. For each genotype, the top three trajectories are shown, ranked by concordance between pseudo-time and true developmental time (DPI). Cells are colored by pseudo-time from early (dark purple) to late (yellow). Related to Figure 7.

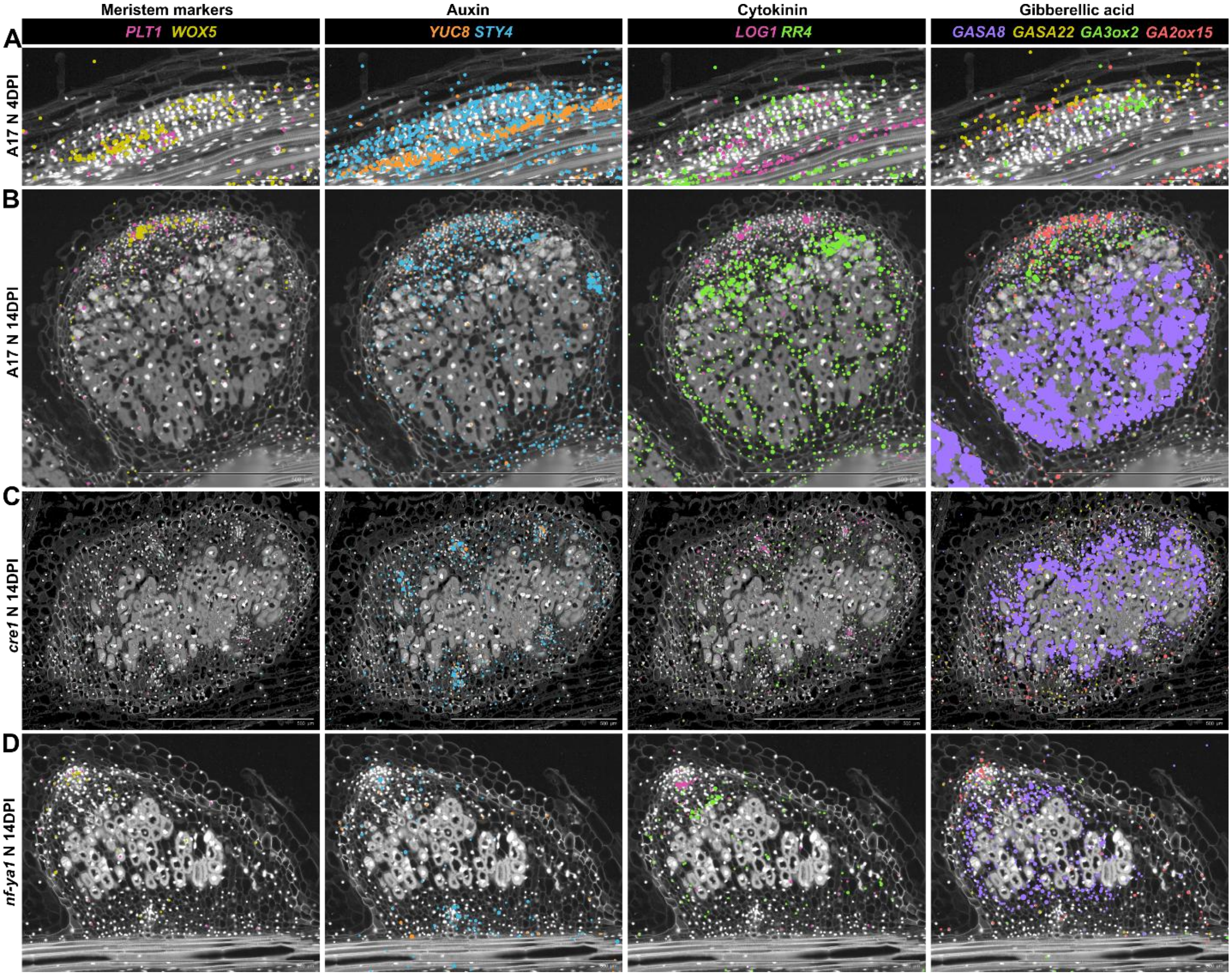


**Figure S13. Spatial coordination of meristem identity and hormone signaling during nodule development**

**(A–D)** Spatial transcriptomic visualization of meristem identity markers and hormone biosynthesis and response genes in wild-type and mutant nodules (A17 background). Transcripts are displayed on representative tissue sections and grouped by functional category, as indicated. From left to right in each panel: (i) Meristem identity markers: *PLETHORA 1* (*PLT1*) and *WUSCHEL-RELATED HOMEOBOX 5* (*WOX5*), marking meristematic and stem-cell–associated domains. (ii) Auxin (IAA) biosynthesis and regulation: *YUCCA 8* (*YUC8*) and *STYLISH 4* (*STY4*). (iii) Cytokinin biosynthesis and response: *LONELY GUY 1* (*LOG1*) and *RESPONSE REGULATOR 4* (*RR4*). (iv) Gibberellin (GA) response and metabolism: *GASA8* and *GASA22* (GA-responsive genes), together with *GA3ox2* (GA biosynthesis) and *GA2ox15* (GA inactivation).

(A) Wild-type nodule at 4 DPI, showing early spatial patterning of meristem and hormone-associated gene expression.

(B) Wild-type nodule at 14 DPI, illustrating refinement of spatial domains during nodule maturation.

(C) *cre1* mutant nodule at 14 DPI, showing altered cytokinin-dependent patterning and disrupted spatial coordination of meristem and hormone-associated gene expression.

(D) *nf-ya1* mutant nodule at 14 DPI, showing perturbed organization of hormone-responsive domains and mislocalization of meristem-associated markers.

**Supplemental Information**

***Table S1. Xenium library and sample metadata***

Metadata for all Xenium libraries generated using the *Medicago* 50-gene panel, the 405-gene spatiotemporal atlas panel and Rhizobia 50-gene panel. The table includes panel information (gene panel size and Xenium library ID), sample metadata (genotype, tissue type, developmental stage, section thickness, and ecotype), and Xenium run information (run batch and raw output availability). Samples include *Medicago truncatula* ecotypes A17 and R108 and mutant backgrounds including *nf-ya1*, *lsh1/lsh2*, *cre1*, *noot1/noot2*, and *lbd16*, spanning nodules and lateral roots across developmental stages from 1–49 DPI.

***Video S1. STPT serial-section imaging and 3D reconstruction of a mature Medicago nodule (49 DPI), related to Figure 2.***

The first 15 seconds show the raw STPT serial-section stack of a mature *Medicago truncatula* nodule. Sequential sections reveal the internal organization of the infection zone, interzone, nitrogen-fixation zone, and early senescence zone. Endogenous autofluorescence, likely arising from NAD(P)H and leghemoglobin, highlights metabolically active rhizobia-infected tissues, while strong cell wall-associated autofluorescence resolves vascular organization across sections.

The second 15 seconds show a 3D reconstruction generated from aligned STPT serial sections. Volumetric rendering reveals the spatial continuity and curvature of the vasculature, the three-dimensional topology of symbiotic zones, and gradients in autofluorescence intensity associated with senescence. The reconstruction enables visualization of spatial relationships between vascular and symbiotic tissues that are not readily apparent in individual two-dimensional sections.
